# Genome wide functional screen for calcium transients in *E. coli* identifies decreased membrane potential adaptation to persistent DNA damage

**DOI:** 10.1101/2020.09.01.277871

**Authors:** Rose Luder, Giancarlo N. Bruni, Joel M. Kralj

**Author notes:** Address correspondence to Joel Kralj.

## Abstract

1.

Calcium plays numerous critical roles in signaling and homeostasis in eukaryotic cells. Unlike eukaryotic cells, far less is known about calcium signaling in bacteria, and few genes controlling influx and efflux have been identified. Previous work in *Escherichia coli* showed calcium influx is induced by voltage depolarization, which were enhanced by mechanical stimulation, suggesting a role in bacterial mechanosensation. To identify proteins and pathways affecting calcium handling in bacteria, we designed a live cell screen to monitor calcium dynamics in single cells across a genome wide knockout panel in *E. coli*. The screen measured cells from the Keio collection of knockouts and quantified calcium transients across the population. Overall, we found 143 gene knockouts that decreased calcium transients, and 32 genes knockouts that increased transients. Knockouts involved in energy production and regulation appeared, as expected, as well as knockouts of the voltage sink, the F1Fo-ATPase. Knockouts in exopolysaccharide and outer membrane synthesis showed reduced transients and refined our model of electrophysiology mediated mechanosensation in *E. coli*. Additionally, knockouts annotated in DNA repair had reduced calcium transients and voltage. However, acute DNA damage did not affect voltage, and suggested that only long term adaptation to DNA damage decreased membrane potential and calcium transients. Our work showed a distinct separation between the acute and long term DNA damage responses in bacteria, which has implications for mitochondrial DNA damage in eukaryotes.

**Importance:** All eukaryotic cells use calcium as a critical signaling molecule. There is tantalizing evidence that bacteria also use calcium for cellular signaling, but much less is known about the molecular actors and physiological roles. To identify genes regulating cytoplasmic calcium in *Escherichia coli*, we created a single cell screen for modulators of calcium dynamics. The genes uncovered in this screen helped refine a model for voltage mediated bacterial mechanosensation. Additionally, we were able to more carefully dissect the mechanisms of adaptation to long term DNA damage, which has implications for both bacteria and mitochondria in the face of unrepaired DNA.

## 2. Introduction

Calcium is an important and ubiquitous signaling molecule in eukaryotic cells^1^, but much less is known about its role in bacteria^2,3^. High resolution tools to measure cellular calcium, including fluorescent biosensors, have enabled detailed knowledge of calcium pathways and dynamics at the sub-cellular level in eukaryotic cells^4,5^, and have recently been applied in bacteria^6^. In bacteria, intracellular calcium concentration changes have been associated with infection^7^, cell division^8^, motility^9,10^, along with other important cellular processes^3^. Bacteria are known to tightly regulate the cytoplasmic calcium levels^11,12^, while changes in the chemical or mechanical environment have been shown to induce calcium transients^6,13–15^. Given the overwhelming importance of calcium in eukaryotes, as well as the many potential bacterial signals, it is critical to understand exactly how and why bacteria regulate calcium levels, as well as the proteins that bind calcium and exert calcium-responsive cellular functions.

One main challenge in uncovering the spectrum of signals encoded by calcium dynamics in bacteria has been a lack of tools to monitor calcium concentration in bacteria with high resolution. Luminescent tools have been used to monitor populations of cells with low time resolution^11^, but they lack the sensitivity to investigate individual cells. The advent of ultrasensitive genetically encoded calcium sensors^4^ was recently applied to *E. coli* and revealed these bacteria act as electrically excitable cells where a voltage depolarization leads to a transient calcium influx, which was tied to mechanical stimulation^6^. These new fluorescent sensors can bridge the single cell gap and potentially reveal new facets of calcium signaling in bacterial cells.

Recently, our group used genetically encoded fluorescent sensors to establish a link between mechanosensation, voltage depolarization, and calcium influx^6^. Similar to vertebrate sensory neurons, mechanically stimulated *E. coli* induced a voltage depolarization, which in turn caused an increase in calcium influx into the cytoplasm. Thus, there is a direct link between mechanosensation and cytoplasmic calcium concentration. Furthermore, mechano- and surface-sensing by bacteria is a critical step in processes associated with human health such as host infection^16^ and biofilm formation^17,18^.

Given the importance of calcium in eukaryotic cells along with the implications in human health, we sought to understand more about the proteins involved in regulating bacterial cytoplasmic calcium concentration. In this paper, we created a genome wide screen of mechanically induced calcium transients in a library of knockouts in *E. coli.* The screen could detect knockouts that decreased or increased the calcium transients. Many hits corresponded with proteins known to influence calcium concentration including genes associated with voltage generation through the electron transport chain and ATP generation through the F1Fo-ATPase. However, we also discovered several pathways that were strongly enriched including outer membrane lipid synthesis and DNA repair. The outer-membrane synthesis knockouts had voltage levels higher than wildtype which suggested a model of mechanosensation in which the chemical structure of the outer-membrane plays a critical role in relaying mechanical signals. Knockouts in the DNA damage repair pathway showed increased DNA damage, but also lower voltage. However, acute DNA damage induced chemically did not change the voltage or calcium transients, which showed that long term adaptation to survive DNA damage included lowering the membrane voltage and growth rates. Though we did not definitely identify a single mechanosensor or voltage gated calcium channel, several hits could potentially provide these elusive functions. Overall, our screening strategy revealed several pathways important in voltage and calcium homeostasis and provided several intriguing hits to follow in future work.

## 3. Results

### High-throughput screen for calcium effectors in *E. coli*

Our model of *E. coli* calcium transients suggested that mechanical stimulation induced voltage depolarization, which led to a transient calcium influx^6^ (Fig 1A). Therefore, a screen for modulators of calcium transients would report conditions that altered mechanosensation, membrane voltage, or calcium handling proteins. To identify genes that modulate mechanically induced calcium transients in *E. coli*, we developed a live cell screen (Fig 1B) using automated 96 well plate imaging of cells expressing a fusion of GCaMP6f (calcium sensor) and mScarlet (insensitive control). The Keio collection of knockouts was transformed with the GCaMP6-mScarlet plasmid and grown on dual selection plates to maintain the gene knockout (kanamycin) and the sensor plasmid (carbenicillin). After transformation, ∼97% of knockouts grew on double selection plates which were moved forward for imaging. Cells were spotted onto agarose pads and pressed into glass bottom well plates (Supp fig 1). Thus, within each well were many cells, each of which had a genotype defined by the Keio collection.

**Figure 1:**
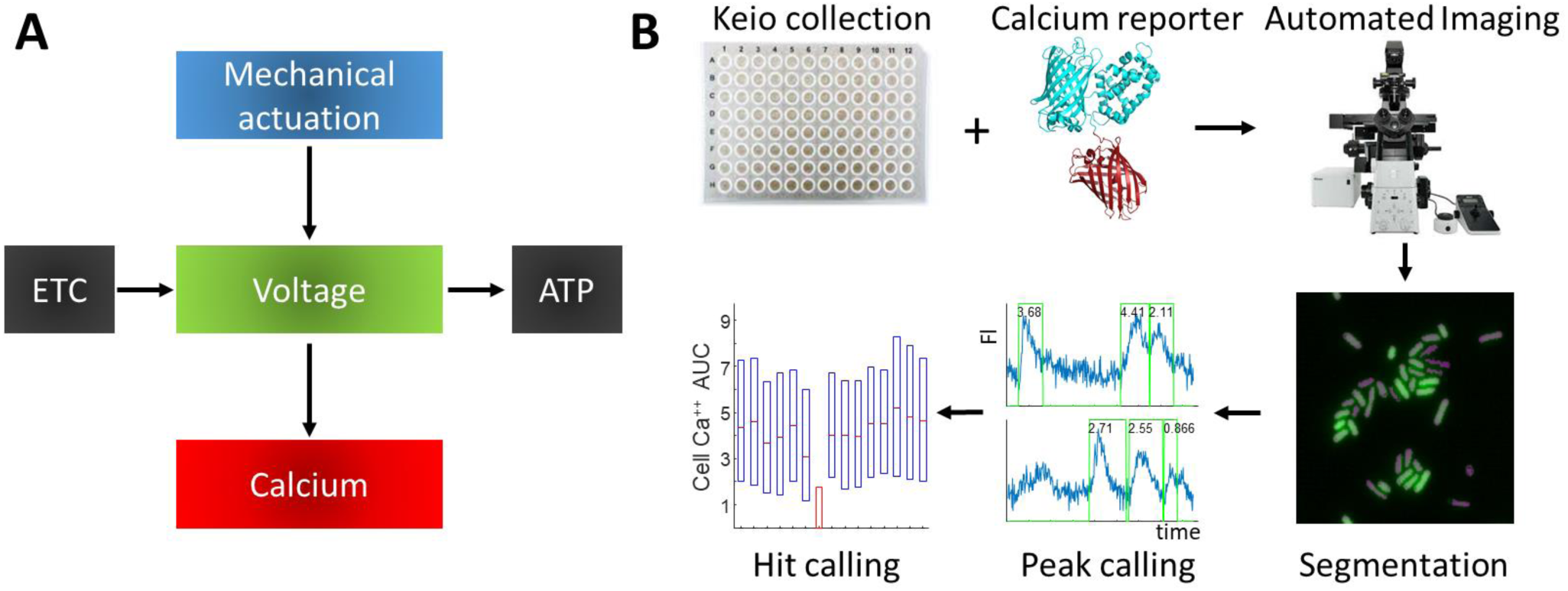
Design for a genome wide screen for calcium effectors in bacteria. (A) Current model underlying the observed calcium transients in *E. coli*. Mechanical stimulation induces voltage depolarizations, which induce calcium influx into the cell. Voltage is generated by respiration via the electron transport chain, and is consumed by the F1Fo-ATPase. (B) Outline of the live cell screen. The Keio collection is transformed with a constitutive GCaMP-mScarlet expressing plasmid and imaged on an automated microscope. Custom analyses segment out individual cells and calculate the calcium AUC. All the cells from a given genotype are then combined, giving rise to the AUC score, which is then screened for knockouts that increase or decrease calcium transients.

From each well, we acquired a 90 second video of the mechanically induced calcium activity to identify genetic knockouts with altered calcium handling. From each field of view (FOV), individual cells were segmented and the mean GCaMP and mScarlet fluorescence was extracted from each frame (see methods, Supp fig 1). From each cell’s fluorescence, we defined transients and calculated the total calcium area under the curve (AUC). Combining every cell from a given genetic identity we assembled a population level AUC as the median of the sum from all cells within a given field of view. Comparing the median population AUC from a given genotype, we were able to identify knockouts that reduced or enhanced the number of Ca^++^ transients (Fig 1B).

To test the resolving power of the screen, CCCP and apramycin were used as positive controls, respectively. These chemical treatments had Z’ factors of 0.32 for CCCP (low transients^6^), and a Z’ factor of 0.28 for apramycin, an aminoglycoside that induces increased transients^19^ (Fig 2A, Supp fig 2). Though the Z’ score was < 0.5, we considered it sufficient to move forward into a genome wide screen by accepting potential false positives during the primary data collection.

**Figure 2:**
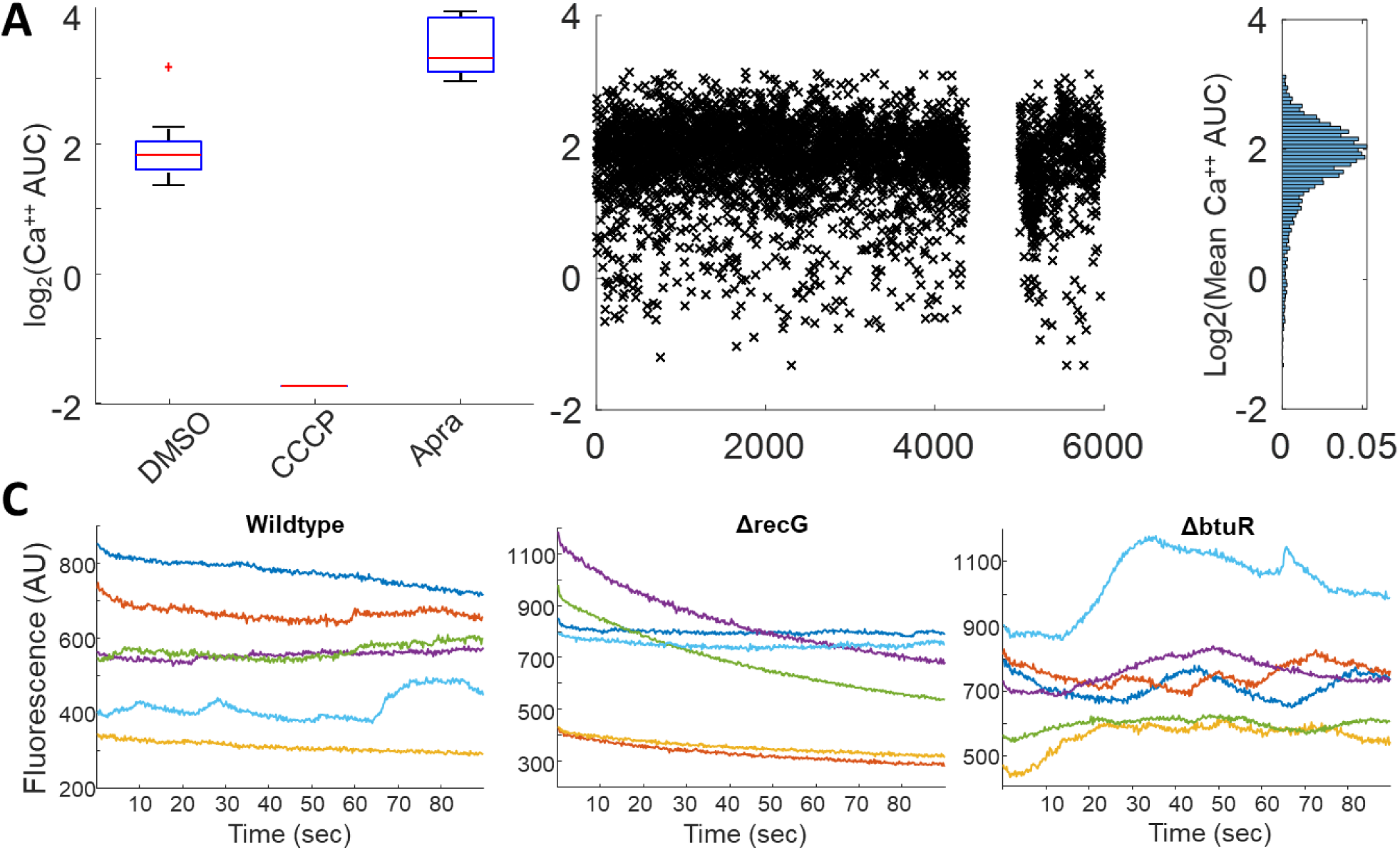
The live cell screen can identify knockouts with altered calcium flux. (A) Box plot of the calcium AUC from cells treated with DMSO (negative control), CCCP (eliminates transients) or apramycin (increases transients). The box plots each have 16 biological replicates. ^*^p < 0.001. (B) Calcium AUC from the primary screen. Each gene in the Keio is plotted on the y-axis, and the log of the calcium AUC is plotted on the y axis. The genes from 4300-5400 are not present in the Keio collection. Right: a histogram of the calcium AUC from the entire screen. (C) Random sample of individual cells from wildtype cells (left), ΔrecG (middle, reduced calcium AUC), or ΔbtuR (right, increased calcium AUC). For all box plots: red line = median, blue box = 25/75 limits, black line = 10/90 limits, red x are outliers.

We then proceeded to use our assay to screen the Keio collection, a genome wide knockout library in *E. coli*^20^. The Keio collection consisted of 45x 96 well plates. We screened all 45 plates as a primary screen, which consisted of raw video data of 7.9 TB. After removing unconfirmed strains, wells too dim to image, and any other artifacts we could determine from the fluorescence images, we extracted time traces from a total of 4.04M individual cells from 3504 genetic knockouts. The top hits from the primary screen were re-imaged in biological triplicate to confirm true positive hits. Across the entire screen, we generated criteria to call knockouts with high and low population wide calcium AUC. Overall, we found 143 knockouts that decreased calcium AUC, and 32 genes that increased calcium AUC (Tables S1,S2). A random sample of traces from ΔrecG (low transients) and ΔbtuR (high traces) qualitatively matched the expectations from the median AUC screen measurements (Fig 2C) when compared to wildtype (WT).

### Expected knockouts in the electron transport chain affect calcium transients

Given our previous work linking membrane voltage with calcium transients, we expected that knockouts known to reduce membrane potential^21,22^ would have a corresponding reduction of observed calcium transients. Indeed, many knockouts of genes in the electron transport chain showed reduced calcium AUC as compared to WT (Table S1). Gene ontology (GO) enrichment analysis^23,24^ of knockouts with reduced calcium transients also showed high enrichment in molecular functions associated with energy production including cytochromes and the TCA cycle. This data provided confirmation that our screen could identify known pathways affecting voltage regulation, which in turn affected voltage transients^6,25^.

In addition to genes involved with voltage generation, we observed that gene knockouts of several components of the F1Fo-ATPase had higher calcium AUC as compared to WT. Considering our current model, these data are also in agreement with the F1Fo-ATPase as a sink of membrane voltage. We expected that inhibiting or eliminating that pathway would result in a higher overall membrane potential and increased calcium transients^15^. Measurements with TMRM confirmed the higher membrane potential as well (Supp Fig 3). Though it has long been known that the electron transport chain generates a voltage that is used by the F1Fo-ATPase^26^, these hits gave confidence that our screen could detect genetic knockouts that affected both voltage generation and mechanically induced calcium transients.

### Outermembrane integrity and exopolysaccharides are critical for mechanosensation

In addition to components regulating voltage, we also expected to find knockouts that affect bacterial mechanosensation. Several inner-membrane (IM) proteins with domains of unknown function were hits with reduced calcium transients, which may be the channels or sensors involved in relaying the mechanical signal across the plasma membrane (Table S1). However, no single IM protein eliminated all calcium transients, which could be due to functional redundancy or a non-proteinaceous calcium channel^27^. However, we did see an enrichment in genes associated with outer-membrane and polysaccharide synthesis that had reduced calcium AUC (Fig 3A). We hypothesized that a fully intact outer-membrane and the presence of polysaccharides are critical components of the mechanical relay. Earlier work showed these knockouts had no reduction in growth rate as compared to WT cells^28^. Furthermore, these knockouts showed no reduction in basal voltage as measured by TMRM when compared to WT cells (Fig 3B), which suggested these knockouts were a part of the mechanosensitive relay.

**Figure 3:**
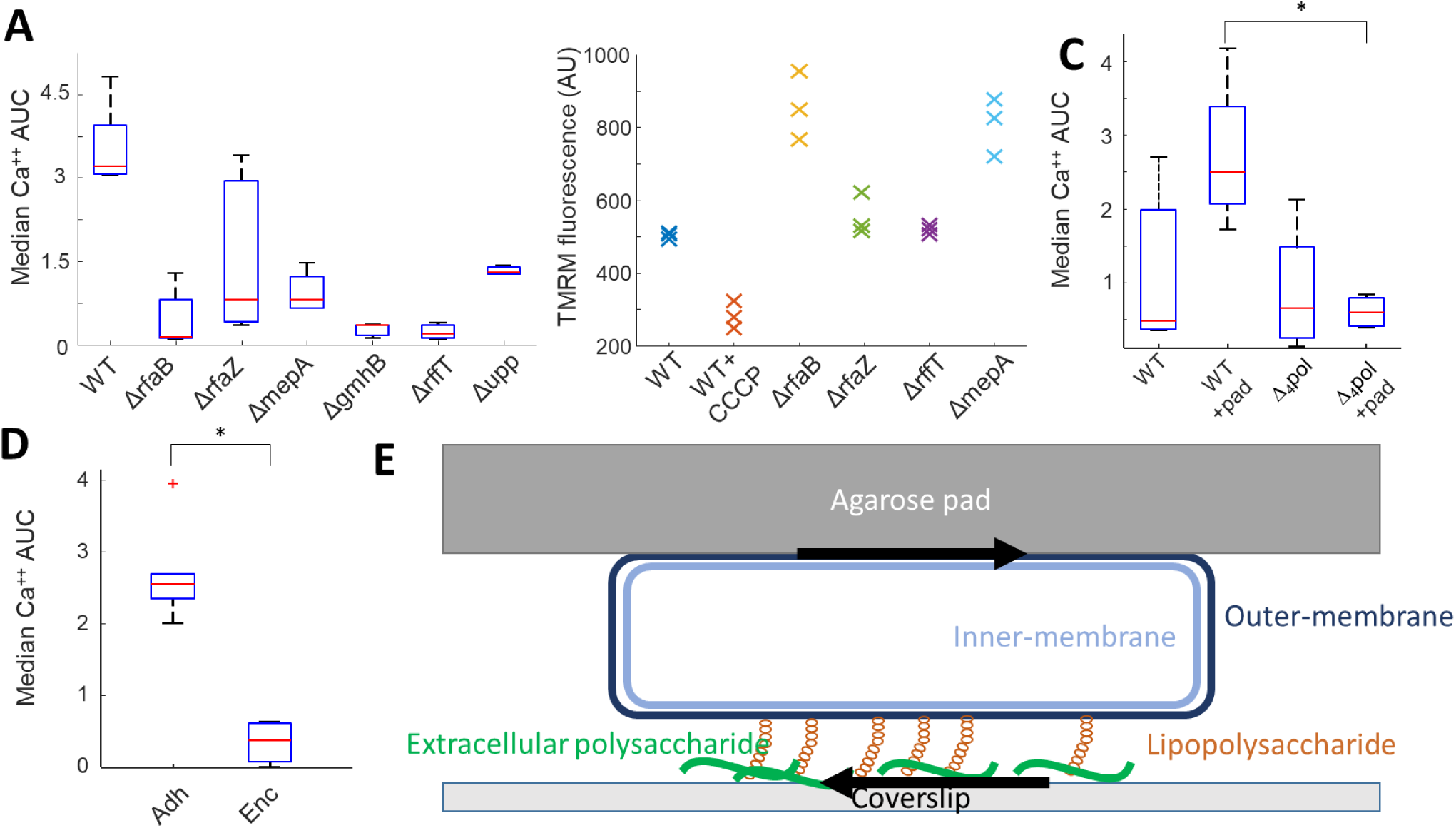
Knockouts of outer-membrane synthesis and EPS secretion refine model for mechanosensation. (A) Box plot of knockouts involved in OM synthesis and EPS secretion had reduced calcium AUC as compared to WT cells. Each boxplot is comprised of 5 biological replicates. (B) Knockouts of the OM synthesis and EPS secretion did not have a reduced membrane potential compared to WT cells. Each genotype was measured in biological triplicate. (C) Box plot of calcium AUC of wildtype cells adhered with poly-l-lysine or an agarose pad compared to a Δ_4_pol knockout under the same conditions. Each box plot is comprised of 4 biological replicates. *represents p < 0.01 (D) Box plot of calcium AUC for *E. coli* adhered under agarose similar to other measurements, or encased in agarose by adding them before solidification of the gel. Each boxplot is comprised of 5 biological replicates. *represents p < 0.01 (E) Proposed model for *E. coli* mechanosensation. A force is applied by the agarose pad, which is opposed by an equal and opposite force form the coverslip. The coverslip force arises from adherence with secreted EPS which then interacts with LPS on the outermembrane of the cell. This force is relayed to a mechanosensor in the inner membrane which remains to be identified. For all box plots: red line = median, blue box = 25/75 limits, black line = 10/90 limits, red x are outliers.

Based on this data and the previous work showing mechanically induced calcium transients, we proposed a model of mechanosensation in which *E. coli* use the outer-membrane and polysaccharide adhesion to stick to the surface of the glass. Upon mechanical deformation with the agarose pad, the outer-membrane undergoes a shear stress which induces the activation of the mechanical sensor, followed by voltage and calcium transients. To test this model, we measured a knockout of 4 components of exopolysaccharide synthesis^29^ (Δ_4_pol) which showed very low calcium transients in our assay (Fig 3C). A second experiment measured calcium transients of bacteria either sandwiched between the glass which is standard in this assay, or encased in agarose which solidified around the cells and exerts no shear force. Cells encased in agarose showed significantly lower calcium transients relative to the glass sandwiched cells (Fig 3D), but continued to grow. Combined, these data are consistent with mechanosensation arising from a shear force relayed through the outer-membrane to induce voltage and calcium transients (Fig 3E).

### Knockouts involved in DNA repair reduce voltage and calcium transients

One unexpected pathway that showed dramatically decreased calcium AUC was genes associated with recombination and DNA repair (Fig 4A). These knockouts had extremely low calcium transients compared to WT cells. This striking data suggested a link between persistent DNA damage and membrane voltage in *E. coli*. DNA damage is known to reduce respiration in bacteria via the SOS response, but respiration is restored upon repair of the damage^30^. Furthermore, DNA damage is associated with depolarized membrane potential in eukaryotic mitochondria^31–33^. Changes to the metabolic state could affect membrane voltage, and previous work established the voltage transients require oxygen^25^.

**Figure 4:**
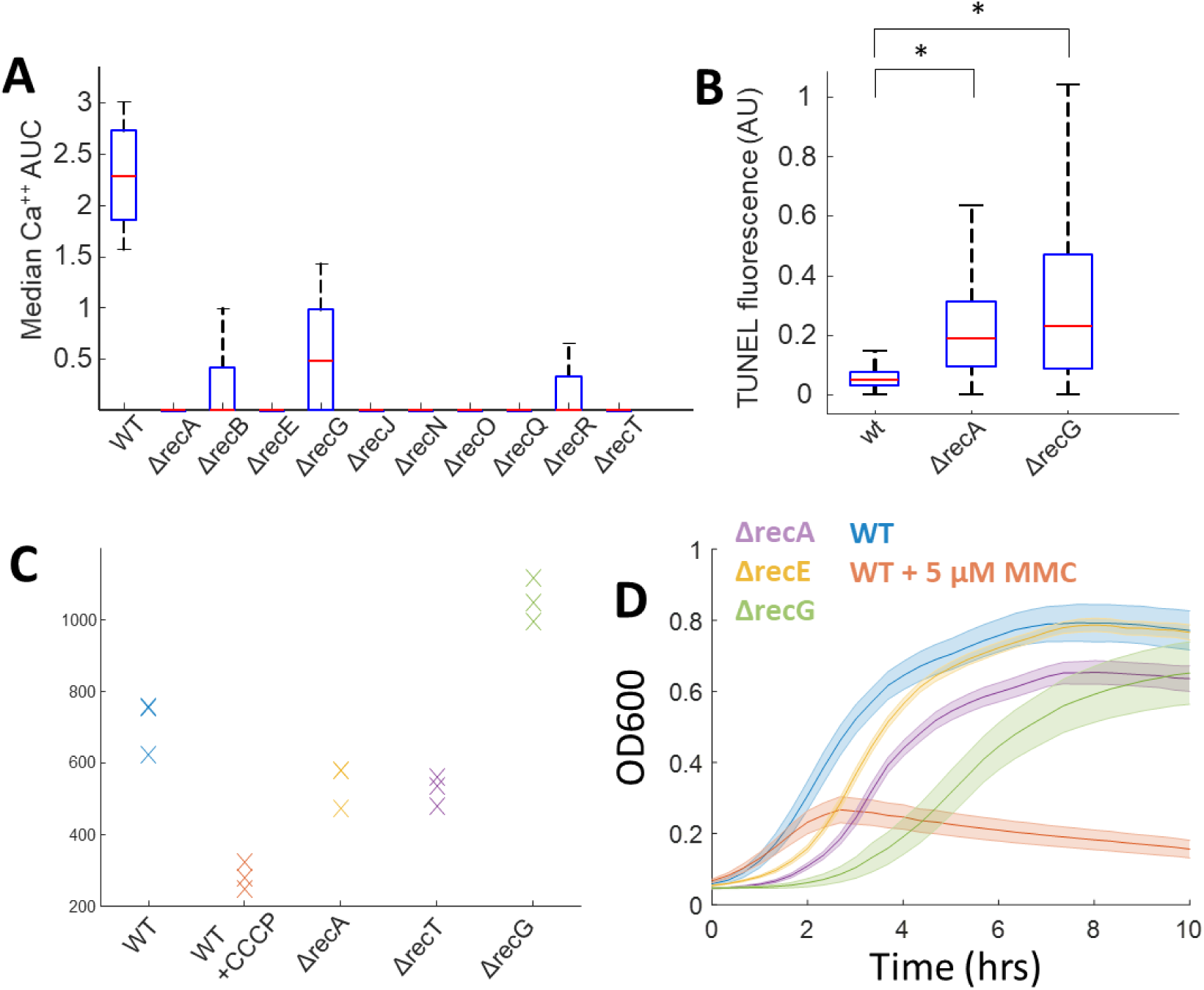
Knockouts of genes associated with DNA repair have reduced calcium AUC. (A) Box plot of knockouts involved in DNA repair compared to WT cells. Each boxplot is comprised of 5 biological replicates. (B) TUNEL fluorescence measured by a flow cytometer for WT, ΔrecA, and ΔrecG. *represents p < 0.05. (C) TMRM fluorescence for comparing WT to rec knockouts. Each marker indicates one biological replicate. Addition of 50 μM CCCP is used to show fluorescence from zero voltage. (D) Growth curves measured by OD600 from WT compared to rec knockout cells. The dark line shows the mean, and the shaded line shows the standard deviation of 3 biological replicates. For all box plots: red line = median, blue box = 25/75 limits, black line = 10/90 limits, red x are outliers.

To first determine if defects in recombination could lead to increased DNA damage, we assayed damage by measuring terminal deoxynucleotidyl transferase dUTP nick labeling (TUNEL). As expected, the knockouts tested showed significantly more DNA damage as compared to WT cells (Fig 4B). Proteins associated with DNA repair are reported to reduce expression of components of the electron transport chain in and mitochondria, so we hypothesized these cells with unrepaired DNA would have lower membrane potential. TMRM measurements showed that recombination knockouts indeed had lower voltage (Fig 4C) as compared to WT cells. The exception was ΔrecG which had a significantly higher TMRM fluorescence than WT cells. We suspected this could be due to cytoplasmic pH differences. For all gene knockouts of DNA repair, there was a corresponding reduction in the growth rate of these cells as compared to WT (Fig 4D). Despite these signatures, the knockout cells observed were not elongated which suggested the SOS response and sulA expression had been reduced^34^. Together, we observed that cells with incomplete repair machinery and persistent DNA damage have reduced voltage and calcium transients, and a reduced growth rate, as compared to wildtype cells.

### Persistent, not acute, adaptation to DNA damage lowers membrane voltage

DNA damage in eukaryotic mitochondria is known to be associated with depolarized membrane potential, and the depolarization has been attributed to ROS buildup and damage to the membrane^35^. In bacteria, it has been suggested the recA filaments interact with the membrane, which could provide a mechanism for reduced voltage^36,37^. The high temporal and spatial resolution of the fluorescent measurements offered the opportunity to investigate the dynamics between DNA damage and depolarization. Genetic knockouts from the Keio collection have many generations of growth and adaptation before imaging is possible, so we instead turned to genotoxic chemicals to induce damage and monitor the acute voltage response of cells.

To control the onset of DNA damage, we used mitomycin C (MMC), a DNA crosslinker purified from *Streptomyces caespitosus*^38,39^. Addition of 5 μM MMC showed bactericidal activity within two hours when measured by CFUs (Fig 5A), confirming high DNA damage within a short time frame. The SOS response was upregulated as measured by filamenting cells 2 hours after treatment (Supp Fig 4). Unexpectedly, calcium transients in cells treated with 5 μM MMC were identical to untreated cells (Fig 5B), and even after 8 hours of treatment, cells showed no difference in calcium AUC as compared to untreated cells (Fig 5C), suggesting the voltage was unchanged. *E. coli* were then grown over 36 hours in liquid culture in the presence of 5 μM MMC, and the cells reached stationary phase which indicated some fraction of the population were adapted to the new environment. These cells, which were continuously treated with the drug MMC, showed severely reduced transients, similar to knockouts of the DNA repair machinery (Fig 5D). Together, our data support a model whereby cells undergo a long-term adaptation to persistent DNA damage by lowering their membrane potential followed by a reduction in the SOS pathway. The adaptation could be through mutation, toxin/antitoxin systems, or others, but they occur at long timescales compared to ROS generation (seconds to minutes).

**Figure 5:**
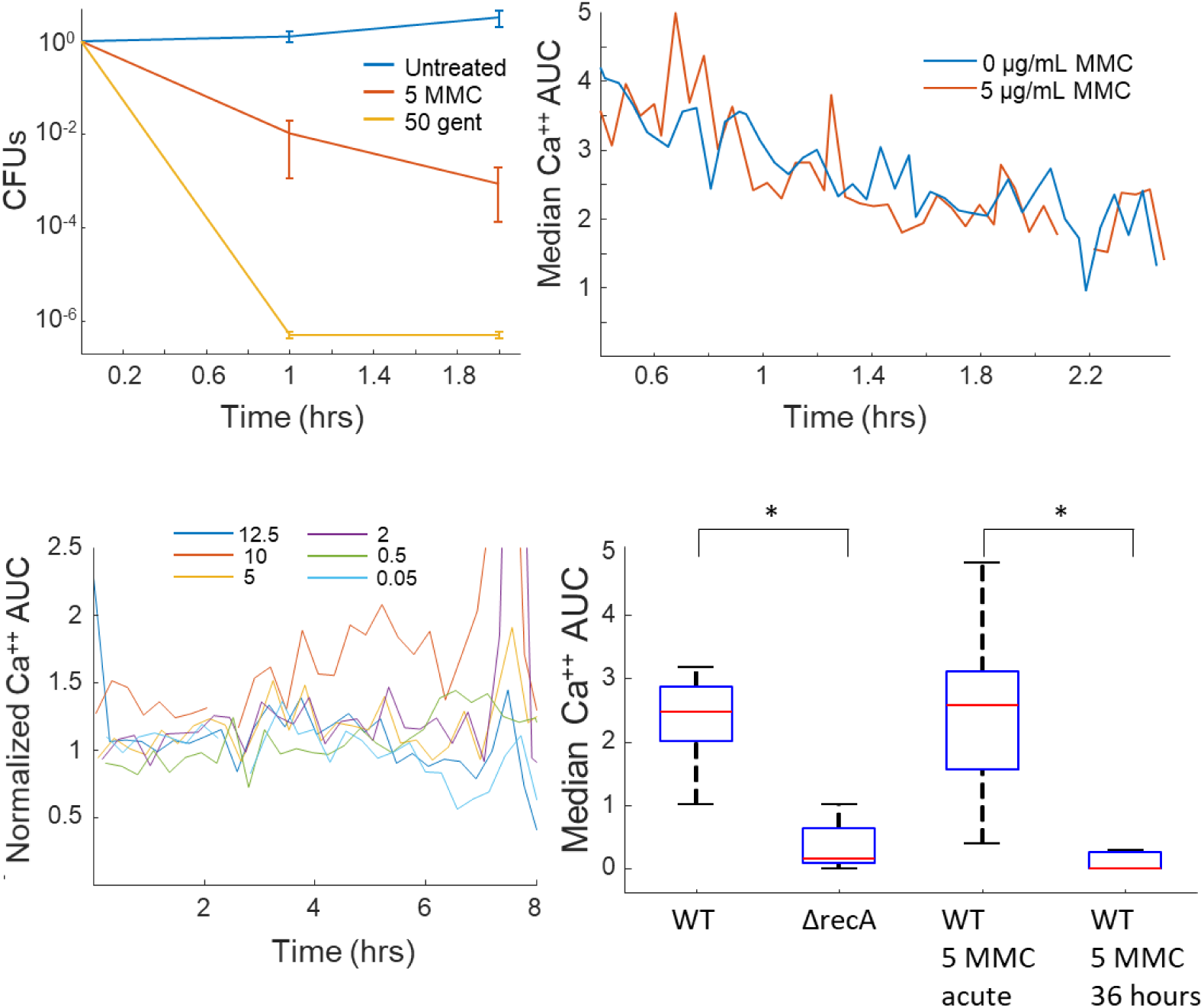
Persistent DNA damage induces an adaption via voltage reduction. (A) CFU assay of cells treated with 5 μM MMC compared to untreated cells. Gentamicin treated cells are a positive control for a bactericidal compound. (B) Time course of calcium AUC taken from 90 sec movies comparing untreated vs 5 μM MMC treated cells. Each point was taken from a unique well, alternating between treated and untreated conditions. (C) Time course of calcium AUC from a titration of MMC concentrations over 8 hours. AUC was calculated from 90 second movies and is normalized to untreated cells. (D) Calcium AUC comparing persistent DNA damage (ΔrecA, 36 hour of 5 μM MMC) to acute DNA damage. * indicates p < 0.01. For all box plots: red line = median, blue box = 25/75 limits, black line = 10/90 limits, red x are outliers.

## 4. Discussion

In this paper, we created a single cell, genome wide screen for modulators of cytoplasmic calcium in *E. coli*. Using this screen, we identified both known and novel pathways that modulate the level of cytoplasmic calcium. The imaging assay had a throughput of 384 genotypes per day, each consisting of dozens to hundreds of single cells, with 450 time points for each measurement. The high signal to noise ratio from the fluorescent signal combined with the automated analytical scripts were able to distinguish positive and negative chemical controls with high confidence. This setup could be easily adapted to other conditions including a chemical screen on calcium AUC, additional bacterial species, or other fluorescent sensors. We believe that single cell, physiological dynamics have the potential to uncover new information and biological processes not available from static imaging.

One of the most interesting and significant hits in our screen was the identification of genes associated with DNA repair, which had reduced voltage and calcium transients as compared to the WT cells. Our data showed the DNA damage clearly results in an impaired voltage, but that the decrease in voltage occurs only over the course of adaption over ∼48 hours. Acute treatment had no impact on the calcium transients, even at concentrations high enough to prevent cells from future cell division. This data suggests a model whereby DNA damage does not immediately lower voltage through ROS production^30^ or interaction with the plasma membrane^36,37^, but rather occurs only as a mutation or transcriptional adaptation that allows a fraction of the cells to continue to divide in spite of persistent DNA damage. Furthermore, the cells turn off the SOS response despite this persistent DNA damage. Understanding the exact mechanisms by which cells accommodate DNA damage via voltage suppression is an important path forward that we plan to study.

DNA damage using MMC also revealed an interesting relationship between cell division as measured by CFUs, and cell metabolism as measured by cytoplasmic calcium maintenance. Addition of 5 μg/mL MMC was enough to prevent > 99% of cells from forming colonies within two hours. Yet, the cells were metabolically active for 8 hours after treatment. This intermediate time period, between treatment and cell death, is similar to previous results with aminoglycosides^19^, and suggests this could be an important consideration during antibiotic treatment if those dying cells can still signal to neighbors, and influence the population level activity.

The mechanisms that tie DNA damage to decreased membrane potential could have impact in the mitochondria of eukaryotic cells as well. Depolarized mitochondria are marked for mitophagy via the PINK-1 pathway, and one trigger is via mitochondrial DNA damage^40,41^. An existing model suggests that DNA damage directly reduces membrane potential through increased ROS^40,42^, but would be the opposite of what we observed in bacteria. One interesting avenue will be to perform similar experiments on eukaryotic cells monitoring the mitochondrial potential upon mitochondrial DNA damage.

## 5. Methods and materials

### Strains and growth

*E. coli* BW25113 was acquired from the Yale Coli Genetic Stock Center and was used as the wildtype strain for all experiments except for MG1655 which was used for comparisons to the Δ_4_pol strain. The Keio collection was purchased from Dharmacon (OEC4988). Keio strains were grown in 10 μg/mL kanamycin to ensure maintenance of the genetic insertion. Cells were grown overnight in LB at 37 °C while shaking at 190 RPM. All strains that carried the GCmS calcium indicator plasmid were grown in 100 µg/mL carbenicillin in addition to other antibiotics necessary to maintain those strains. Δ_4_pol strain was maintained in media with 10 µg/mL chloramphenicol, 30 µg/mL kanamycin, and 25 µg/mL zeocin.

### Plasmids and transformation

The constitutive GCaMP-mScarlet plasmid was generated in a previous publication^19^ and is available on Addgene (#158979). The plasmid was transfected into *E. coli* knockouts by using the Transformation and Storage Solution (TSS) buffer in a 96 well plate. Briefly, cells were grown overnight in a deep well 96 plate in LB and kan. Cells were then spun down at 1900x g for 7 min, and the supernatant was removed by pouring. TSS buffer was added, and incubated with 1 uL per well from a miniprep of the plasmid. Upon incubation for 1 hour, 5 μL of the cells were placed onto a double selection plate (carb – 100 μg/mL, kan – 50 μg/mL) using a multichannel pipet, and the plates were incubated overnight at 37 °C.

Colonies of the transformed cells were then picked using a pin transfer tool, and grown up overnight in LB (carb – 100 μg/mL, kan – 50 μg/mL), and glycerol stocks were made from this suspension. To grow cells for imaging, the glycerol stock in 96 well plates was placed into a 96 well plate with LB (carb – 100 μg/mL, kan – 10 μg/mL) using a pin transfer and cells were grown overnight at 37 °C while shaking.

### TMRM assays

A 1 mL Falcon polystyrene round-bottom tube with 940 µL of M9 minimal media with 0.4% glucose and 1X NEB amino acids was seeded with 50 µL of overnight growth suspension of the strain to be tested (in biological triplicate), and grown at 28°C with 200 rpm shaking. When the cells reached ∼0.4 OD, 10 µL of TMRM was added to the suspension at a final concentration of 0.2 mM TMRM. Thirty minutes later, cells were quantified for their TMRM incorporation by counting 100,000 events per condition using a BDFACSCellesta Flow Cytometer with the following Voltage settings: FSC at 700, SSC at 350, with 561 nm laser D585/15 at 500, C610/20 at 500 and B670/30 at 481. Emission for each event was collected at the 585/15 nm wavelengths.

### Tunel Assay

DNA damage was measured using the TUNEL APO-DIRECT kit (BD 556381) using the manufacturer’s suggested protocol. Cells were fixed with ethanol followed by washing and staining.

### Keio imaging

Imaging took place on an inverted microscope setup using 96 well glass bottom plates (Brooks, MGB096-1-2-LG-L), similar to earlier work^19^. Cells grown in overnight cultures in 96 wells at 37 °C in LB with carbenicillin to maintain the plasmid. Agarose pads in a 96 well format were created using a custom 3D printed mold (https://www.shapeways.com/shops/kraljlab). The mold was designed so that each well held a 200 μL agarose pad. Agarose was dissolved at a final concentration of 2% in partial minimal medium (PMM, 1x M9 salts, 0.2 % glucose, 200 μM MgSO4, 10 μM CaCl_2,_ 1x MEM amino acids) and added into the mold which was covered by a piece of glass (McMaster Carr, 8476K43). A second piece of glass was added to the to create flat surfaces on both sides. After the agarose gel solidified (typically > 30 minutes), 2 μL of the cell suspension was added onto each pad and the solution was allowed to fully diffuse into the gel. The pads were then pressed into the glass bottom plate using a second 3D printed piece. There is an inversion of the plate (A1 from cell suspension -> A12 in the glass bottom plate) that was accounted for in the analysis software.

Imaging took place on an automated inverted microscope (Nikon Ti2) using a 40x NA 0.95 objective to image onto two sCMOS cameras (Hamamatsu, Flash 4 v2) using a custom dichroic splitter in the emission path. Emission filters were a 525/50 (GCaMP) and 568 LP (mScarlet) and we used a 561 LP dichroic to separate the two colors. Cells were illuminated using an LED source (Lumencor, Spectra X), using simultaneous excitation with 470 nm and 550 nm light. Images were captured continuously with an exposure time of 200 ms (5 Hz) for a total of 90 seconds.

To enable automated imaging of all 96 wells without additional user input, a single point was selected in each well by the user such that there were cells, but not too dense. Selecting all 96 points took on average ∼20 minutes. Enabling the Perfect Focus on the Ti2 kept the plate in focus across the entire well plate. Capture of the entire movie took ∼2.5 hours per plate, so we were typically able to image 2-3 plates per day. Image data was stored with the metadata in the .nd2 format for downstream processing.

### Image processing

Image processing from the movie files was similar to our previous reports^19^, with a few changes due to the nature of the movie collection. Image analysis occurred in Matlab using custom .m scripts. Image processing followed the general scheme of (1) estimating the illumination profile for all experiments on a given day, (2) correcting the uneven illumination for each movie, (3) registering drift and jitter in XY, (4) subtracting an estimated background, (5) segmenting cells using a Hessian algorithm, (6) extracting time traces for individual cells, (7) processing each time trace to calculate calcium area under the curve (AUC).

1. Estimating the illumination profile: For a given day, every movie was averaged across time, and opened using a morphological operator and blurred using a 2D Gaussian filter. Each of these experimental images were then averaged together to give an estimate of the uneven illumination. These images were smooth across the entire field of view, and varied by ∼50% across the entire image.
2. Correcting uneven illumination. Each individual movie was then loaded into memory sequentially. Each frame of the movie was converted to a double, and then divided by the uneven illumination. This image was then multiplied by the average value of the movie and converted back into a uint16 to maintain consistent intensity values. Each frame was then reassembled into an illumination corrected movie.
3. Registering drift and jitter in XY: Each frame was aligned to the previous frame using a convolution of the 2D Fourier transform. Each sequential image was first estimated by applying the XY warping from the previous frame. Then, the 2DFT was taken for each image, and multiplied to the previous frame. The optimal updated XY position was then calculated and applied. Due to the short time of these movies (90 sec), little drift was observed for most measurements.
4. Subtracting the estimated background: The background was estimated for each frame individually using a morphological operator. A disk structured element with radius 9 µm was blurred with a Gaussian filter. This background estimation was then subtracted from the original image. To protect against potential negative values, the minimum of the entire movie was set to 50 counts.
5. Segmenting cells using a Hessian algorithm: To segment cells, first the foreground was estimated using Otsu’s method from the background subtracted image. The Hessian was then calculated on the background subtracted image, and then elementwise multiplied to a logical image of the foreground. Otsu’s method was again used on this modified Hessian image to identify individual cells. Hard limits were set to remove potential noise that did not fit given criteria for size or minimum intensity. We found that first increasing the size of the image using a spline interpolation gave superior segmentation results. Using this method, not all cells were identified within a microcolony, though we estimate that it can identify ∼96% of the cells accurately.
6. Extracting time traces for individual cells: From a given identified cell, for each time point in the movie, we extracted the mean intensity using the Matlab command, regionprops. The mean intensity for both the GCaMP6f and the mScarlet were extracted using this method, or any other fluorophore the cells expressed.
7. Processing each time trace to calculate the calcium AUC. To identify calcium transients in a given cell, we first removed any low frequency photobleaching using a moving median filter, followed by a wavelet denoising (wdencmp.m). To identify the transient starts, any point of the differentiatl that rose above zero (rising slope) and enforced that a peak have an area under the curve > 3σ. For any given cell, all the transients that met these criteria were summed to give a cellular AUC over the duration of the movie. To calculate the genotype AUC, the median AUC of the population was calculated to minimize any outliers from non-stationary cells.

### CFUs

CFUs were measured by plating treated cells onto LB-agarose without antibiotic and counting growing colonies. CFU measurements were conducted trying to mimic the experiments performed via microscopy. Briefly, cells were grown overnight in LB and diluted 1:20 in 5 mL PMM. These cultures were grown at room temperature and shaking for 2 hours (t = 0) followed by the addition of antibiotic. At each time point, the culture was removed from the shaker, and 100 µL was removed. A 10x series dilution was then conducted by removing 20 µL and adding to 180 µL LB alone in a 96 well plate. The 10-fold dilution was performed 7 times, leading to the original concentration to a dilution of 10^7^. From each of the 10x dilution series, 3 µL was plated onto an LB agar pad and left to dry (1 colony = 333 cells/mL, lower end of our dynamic range). After an entire experiment (typically 5 hours), the agar was placed into an incubator and grown overnight. Colonies were then manually counted the next morning.

### Growth curves

Growth curves were collected in a 96 well plate using a Tecan Spark plate reader. The plate reader was set to 37 °C and data was collected from each well every 5 minutes over 10 hours.

## Acknowledgments

Special thanks to Theresa Nahreini for help with cytometery. Thanks to Bianca Audrain, Christophe Beloin, and Jean-Marc Ghigo for the Δ_4_pol strain.

## Supplementary figures and tables

**Supplementary Figure 1:**
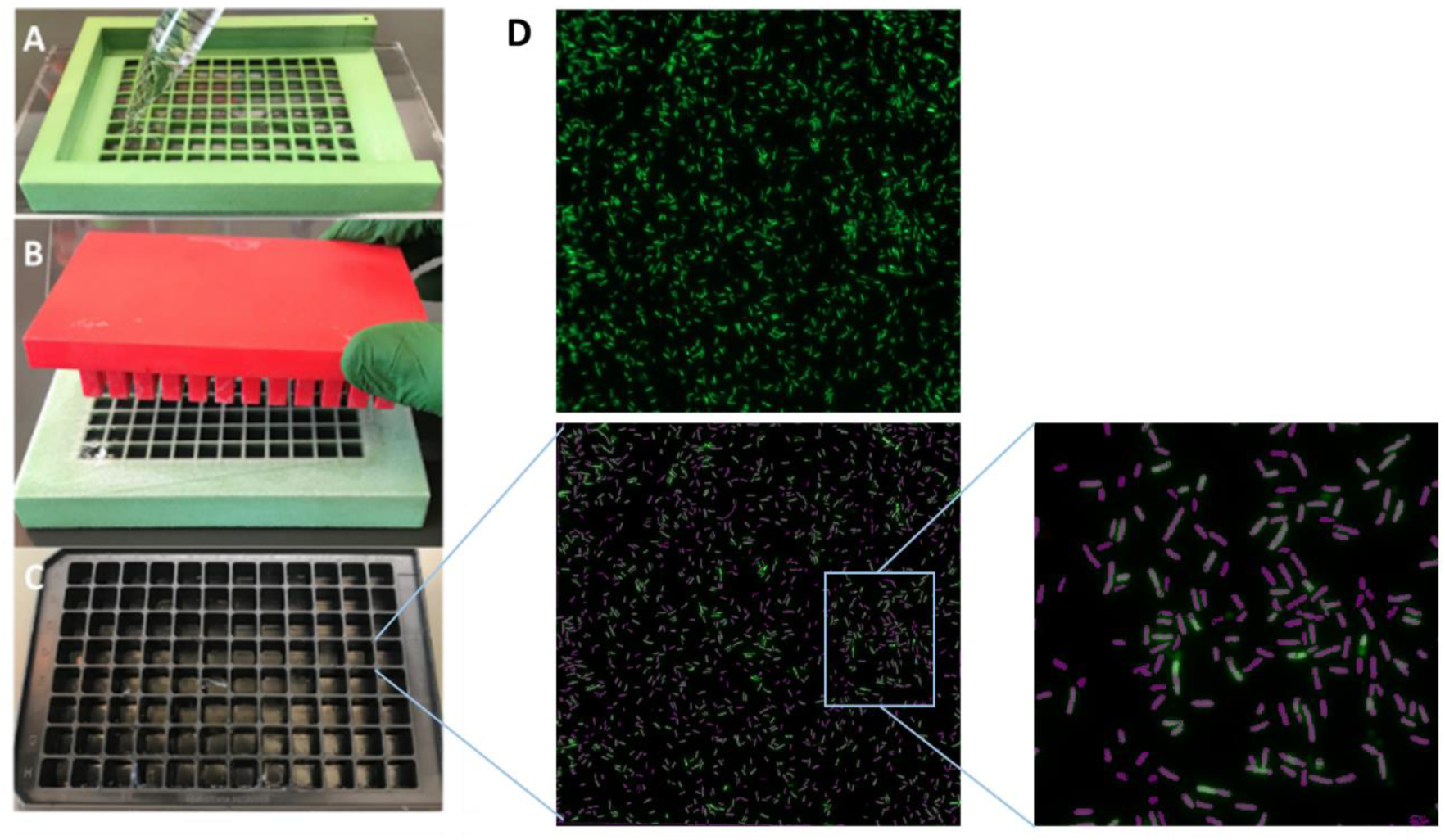
Loading and imaging cells in a 96 well plate. (A) Pictures of the 3D printed agarose mold to immobilize *E. coli* in glass bottom plates. The green mold is filled with agarose between 2 pieces of glass until solidification. Cells are added to the bottom of the pad in 2 μL drops. (B) The red press is used to push each agarose pad into (C) the bottom of the glass bottom dish. (D) Top: 160 × 160 μm^2^ FOV of cells expressing GCaMP mScarlet immobilized under an agarose pad. Bottom: Overlay of the image with the Hessian calculated cell mask shown in purple. The mask calculated 1460 individual cells in this particular FOV. Right: Zoom in on one section of cells and mask.

**Supplementary Figure 2:**
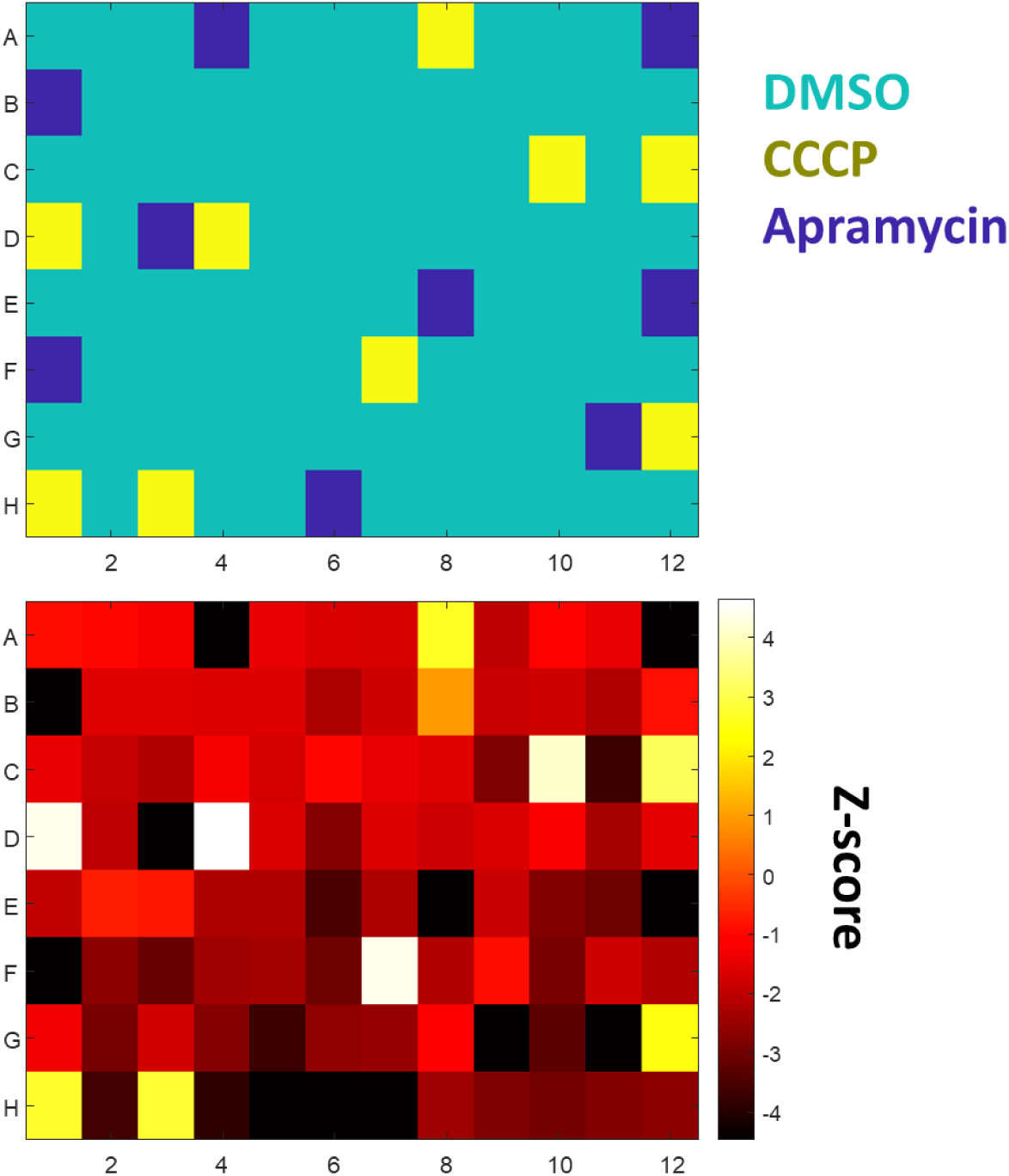
Control plates to test high and low calcium AUC relative to WT cells. (A) A plate map showing cells treated with DMSO (negative control), 50 μM CCCP (positive control, low AUC), or 100 μg/mL apramycin (positive control, high blinks). (B) Measured Z-score from the test plate with the Z-score for each well color coded.

**Supplementary Figure 3:**
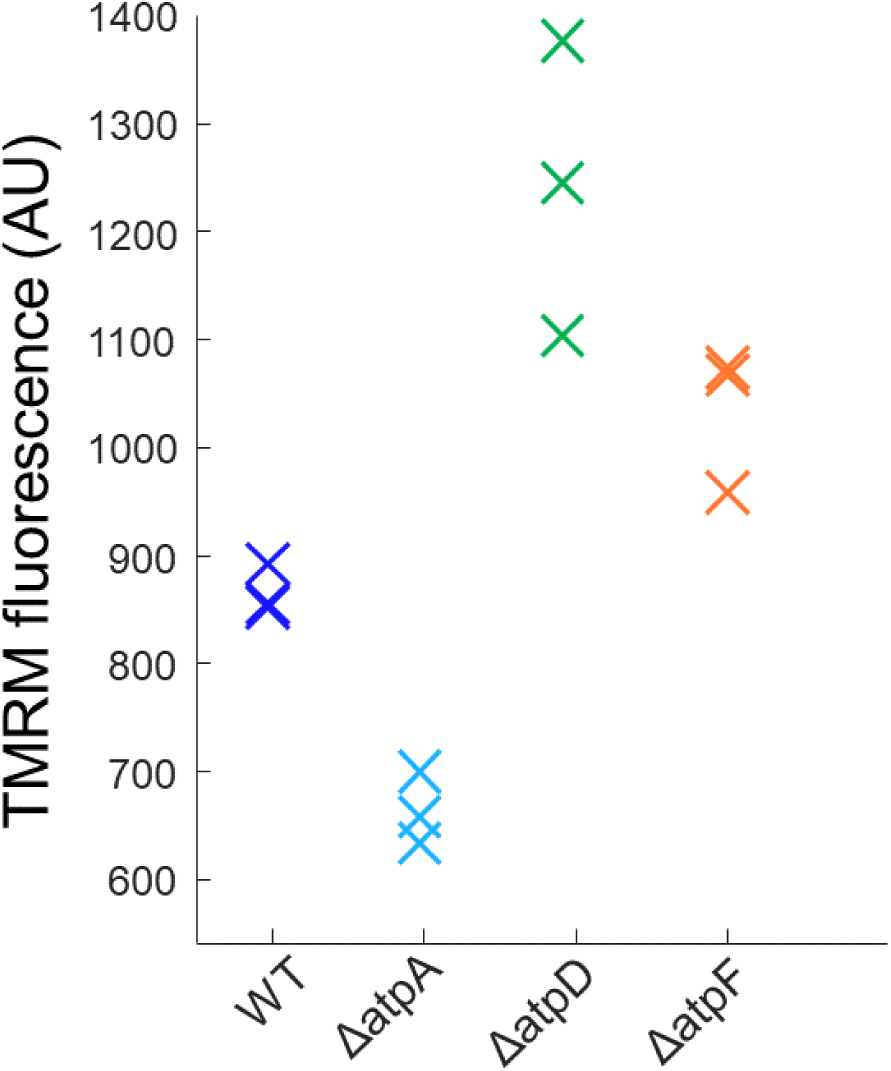
Knockouts of the F1Fo-ATPase had altered membrane voltage that correlated with calcium transients. Each mark is the median of 100k events measured on the cytometer, and each genotype was measured in biological triplicate. ΔatpA had reduced calcium transients and voltage, while ΔatpD and ΔatpF had increased voltage and calcium transients compared to WT.

**Supplementary figure 4:**
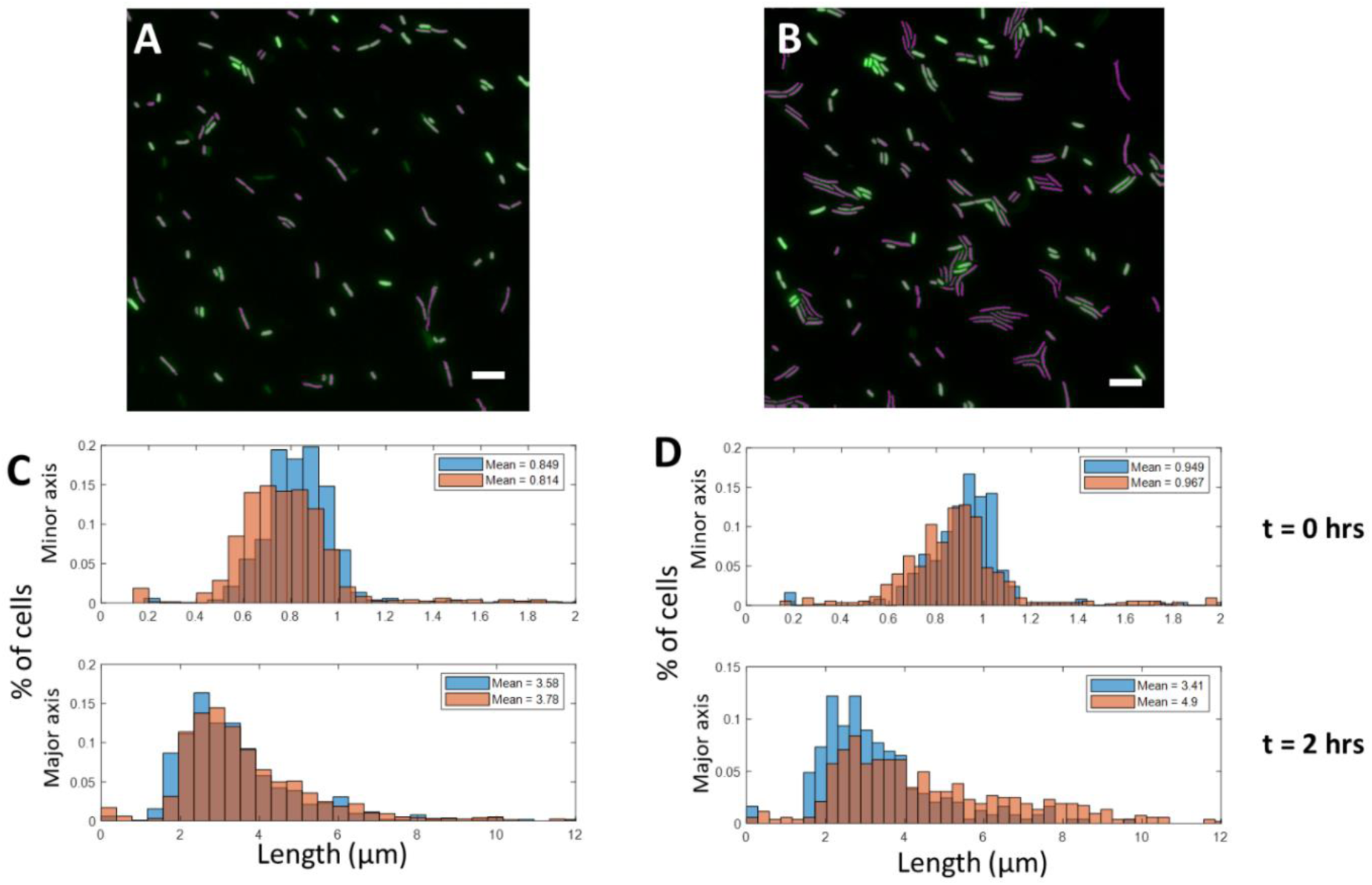
Cell morphology before and after treatment with 5 μM MMC. (A) GCaMP fluorescence (green) and the segmented mask (purple) before addition of drug and (B) 2 hours after drug. (C) Minor axis (orange) and major axis (blue) distributions at t = 0 hours before drug addition calculated by the masks shown in (A). (D) Minor axis (orange) and major axis (blue) distributions at t = 2 hours before drug addition calculated by the masks shown in (B).

**Table S1:**
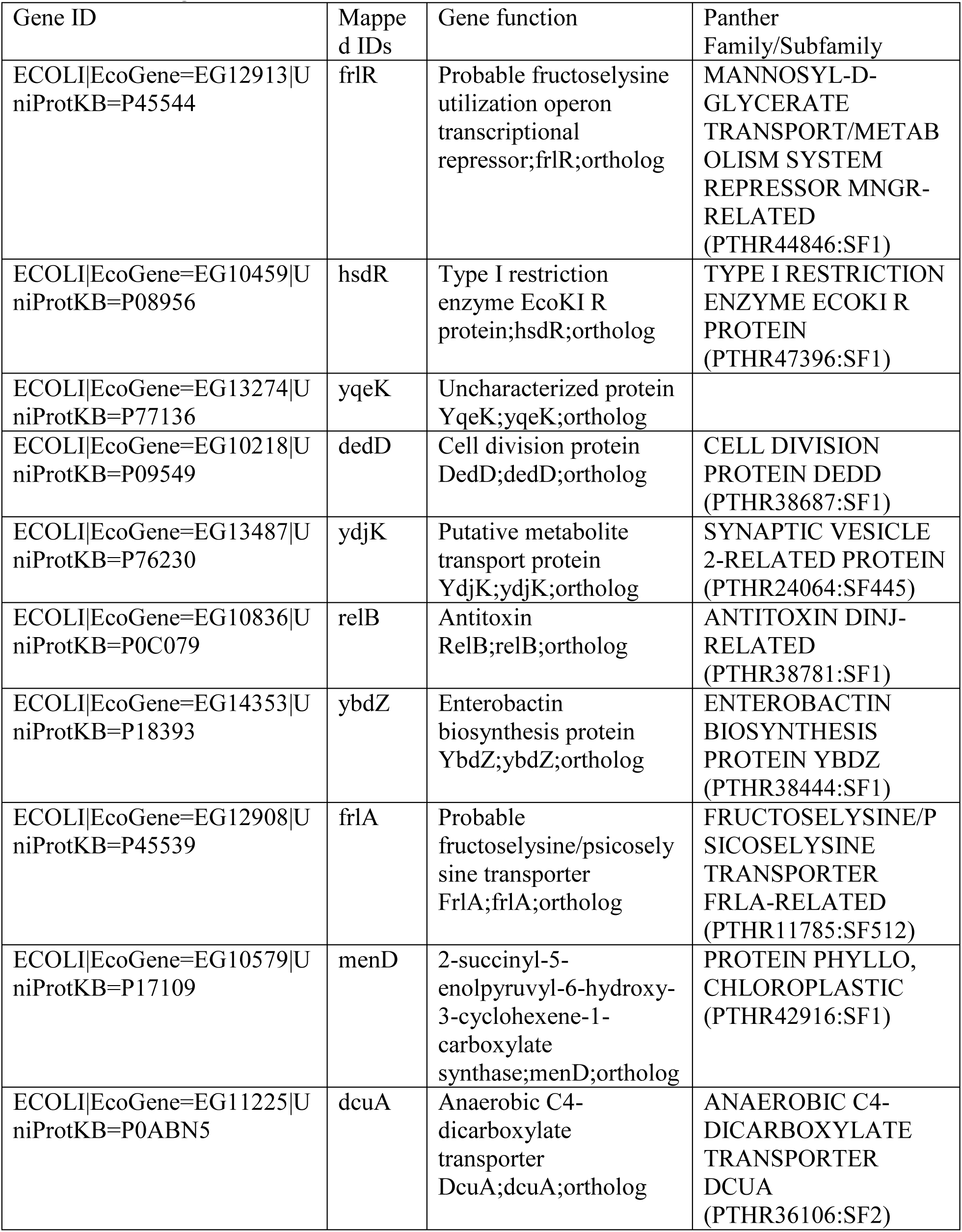
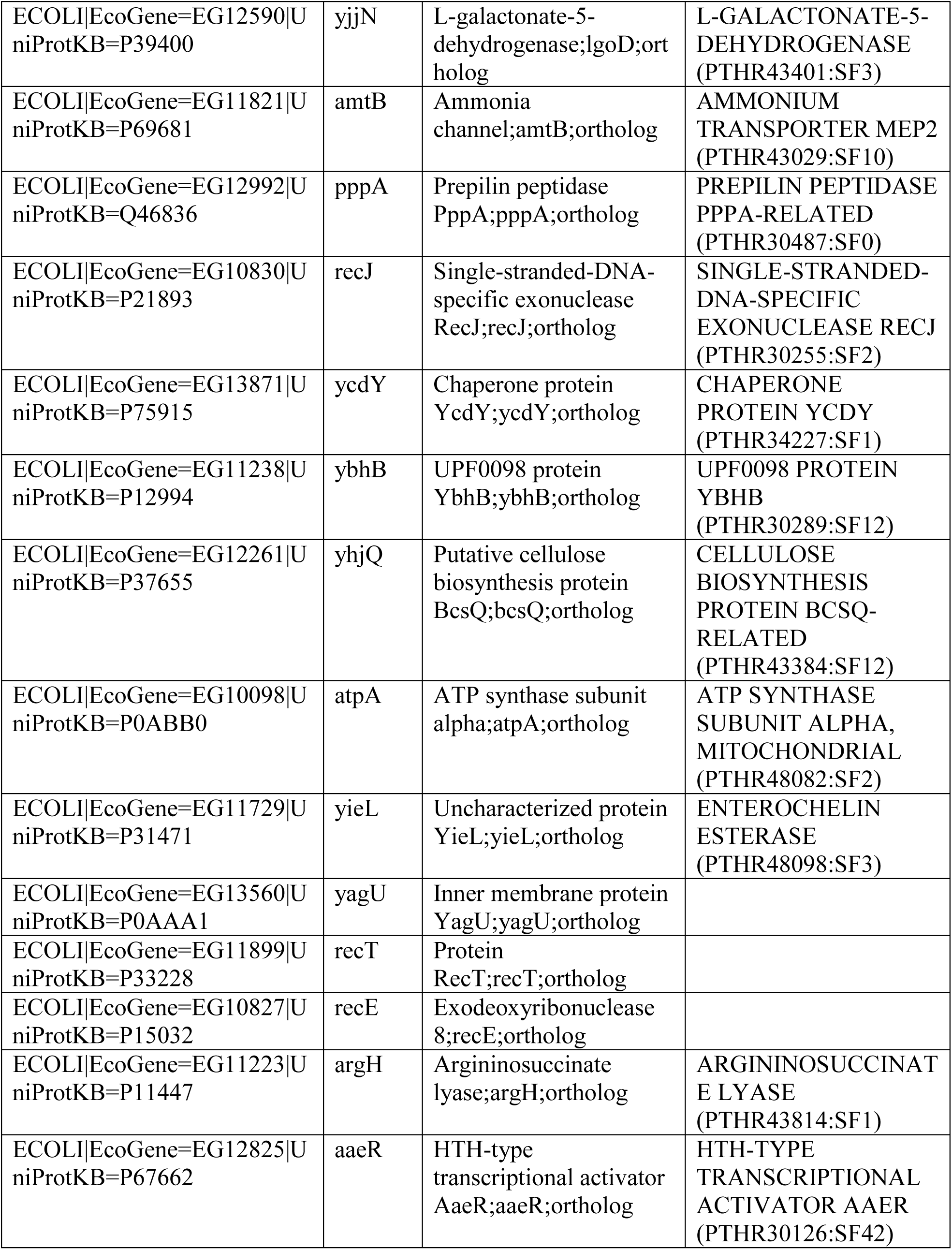
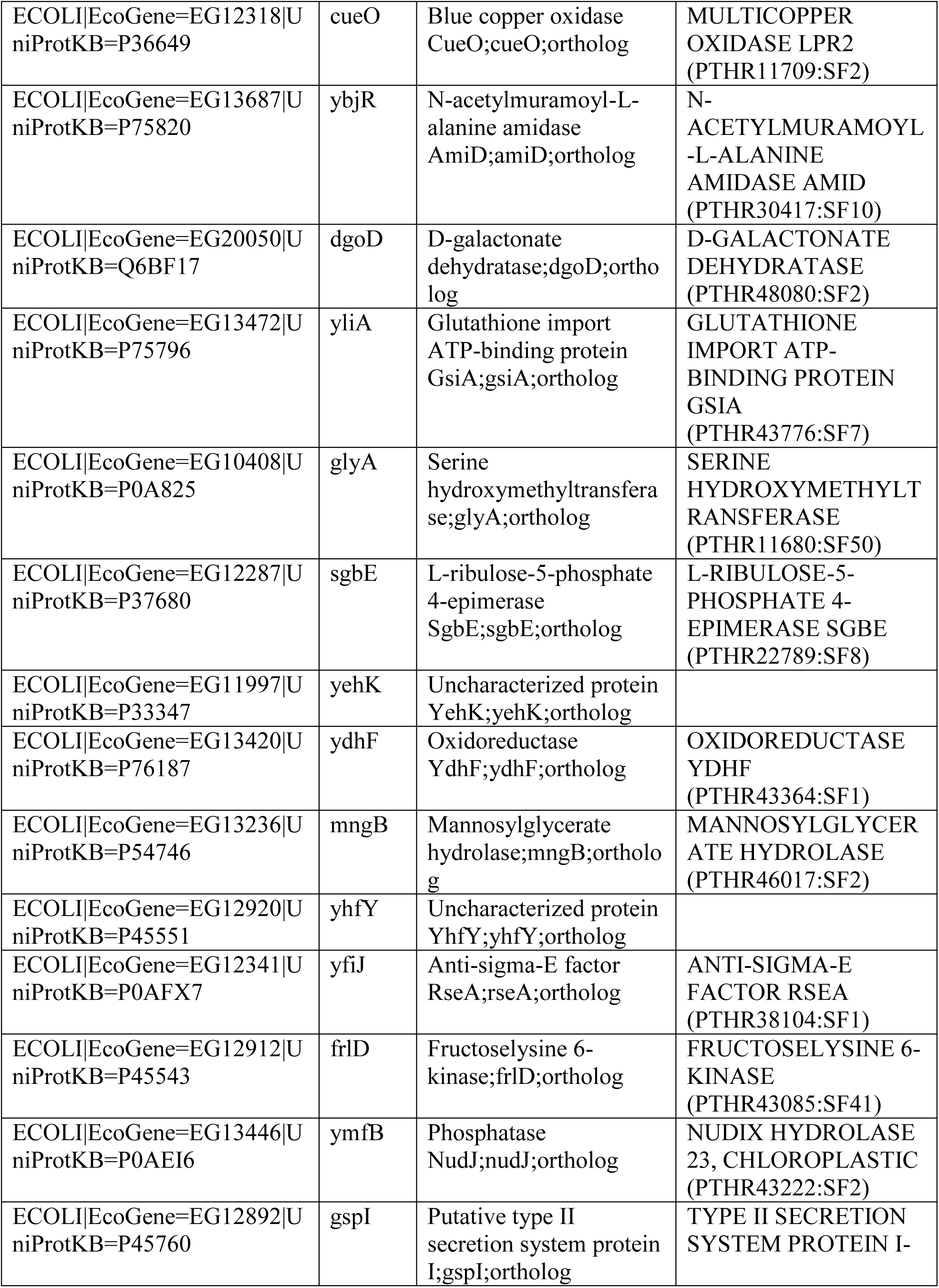
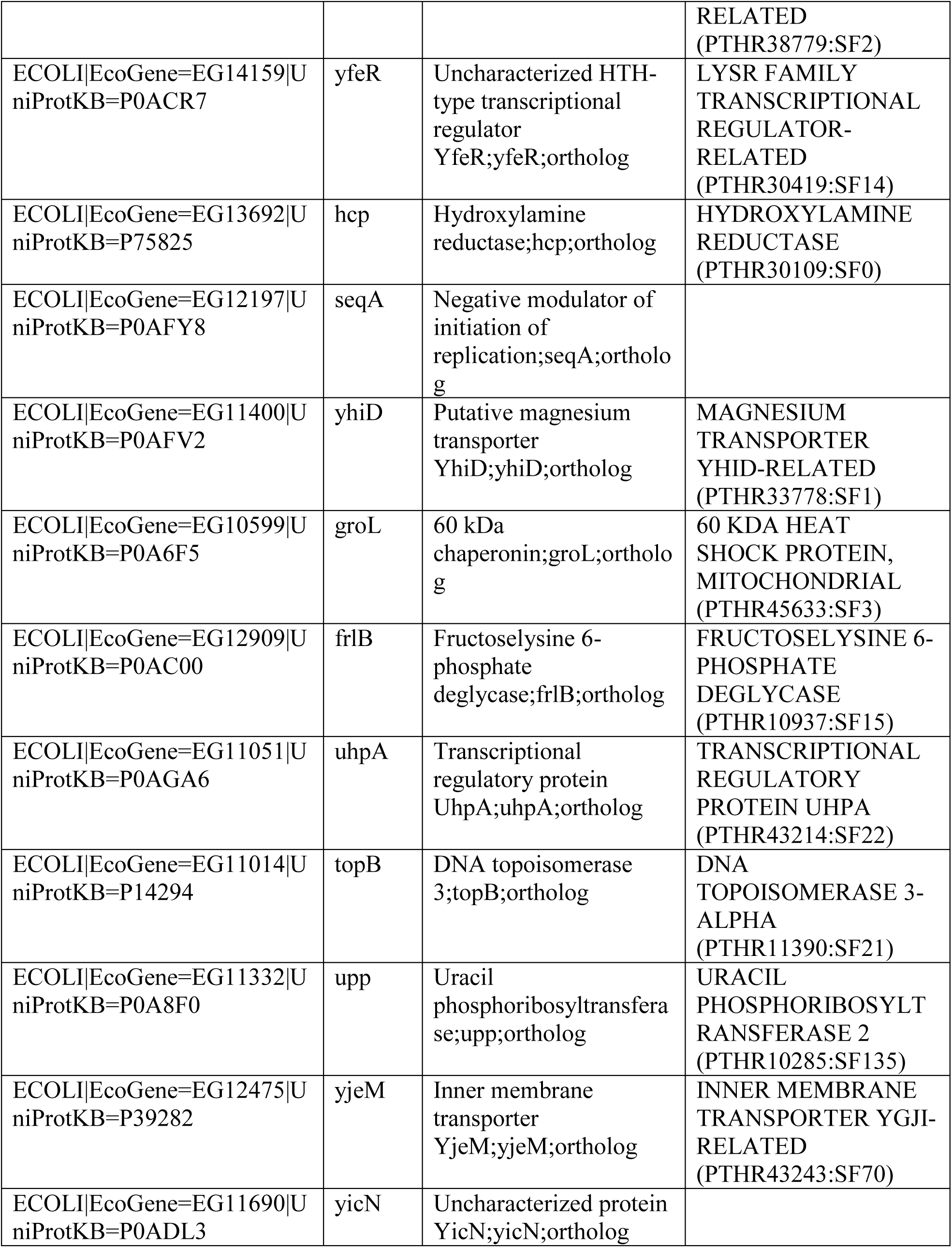
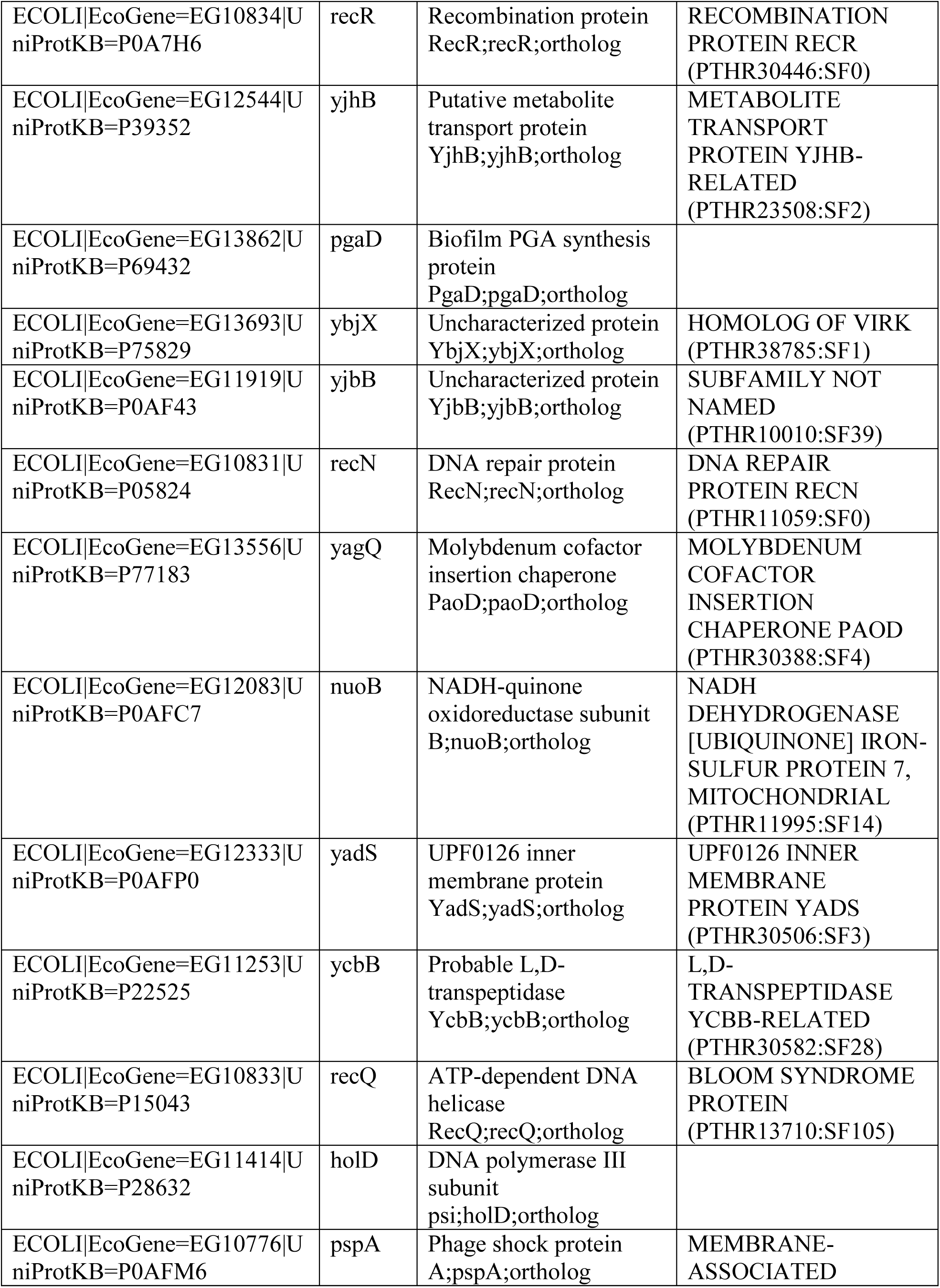
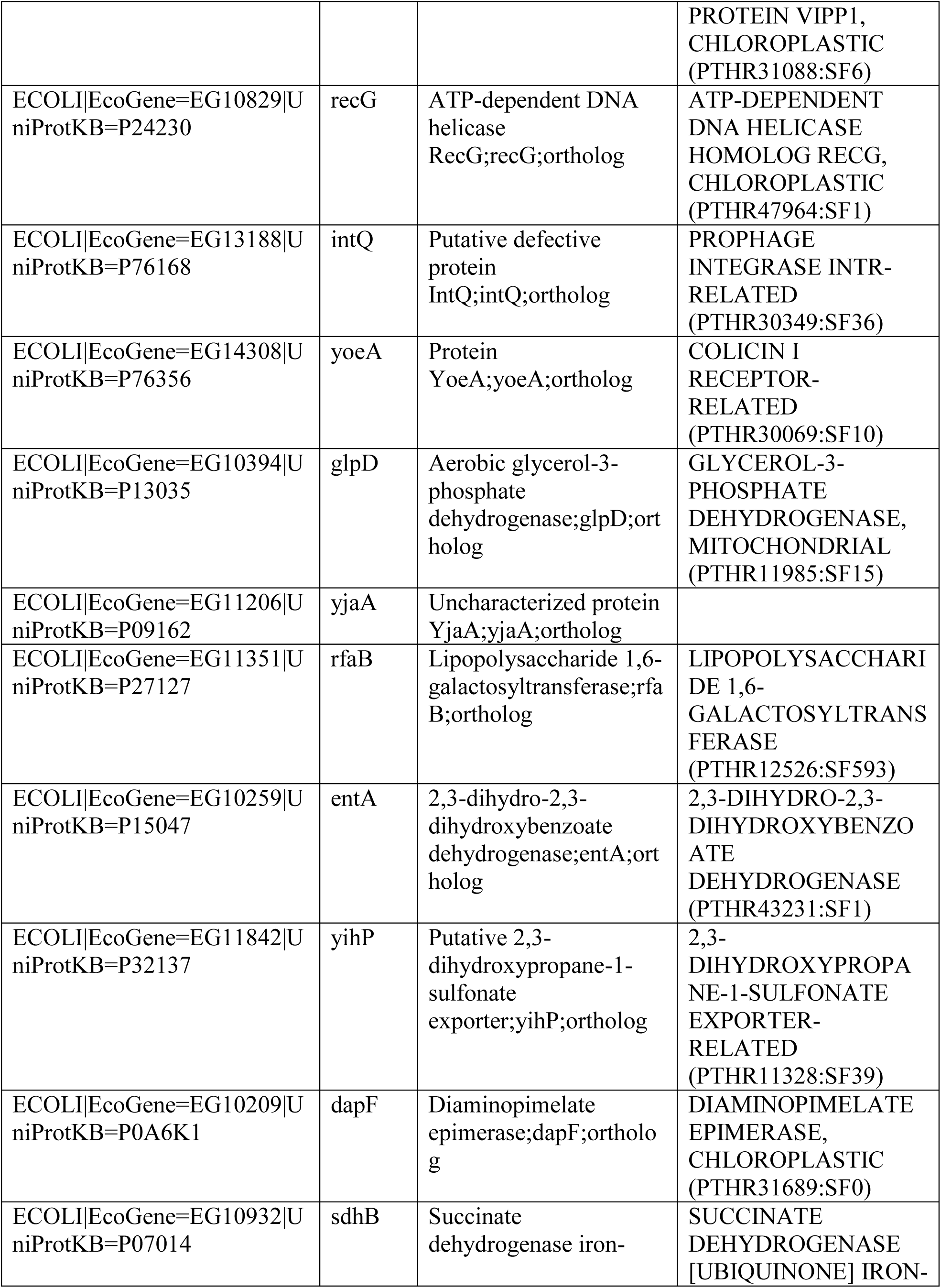
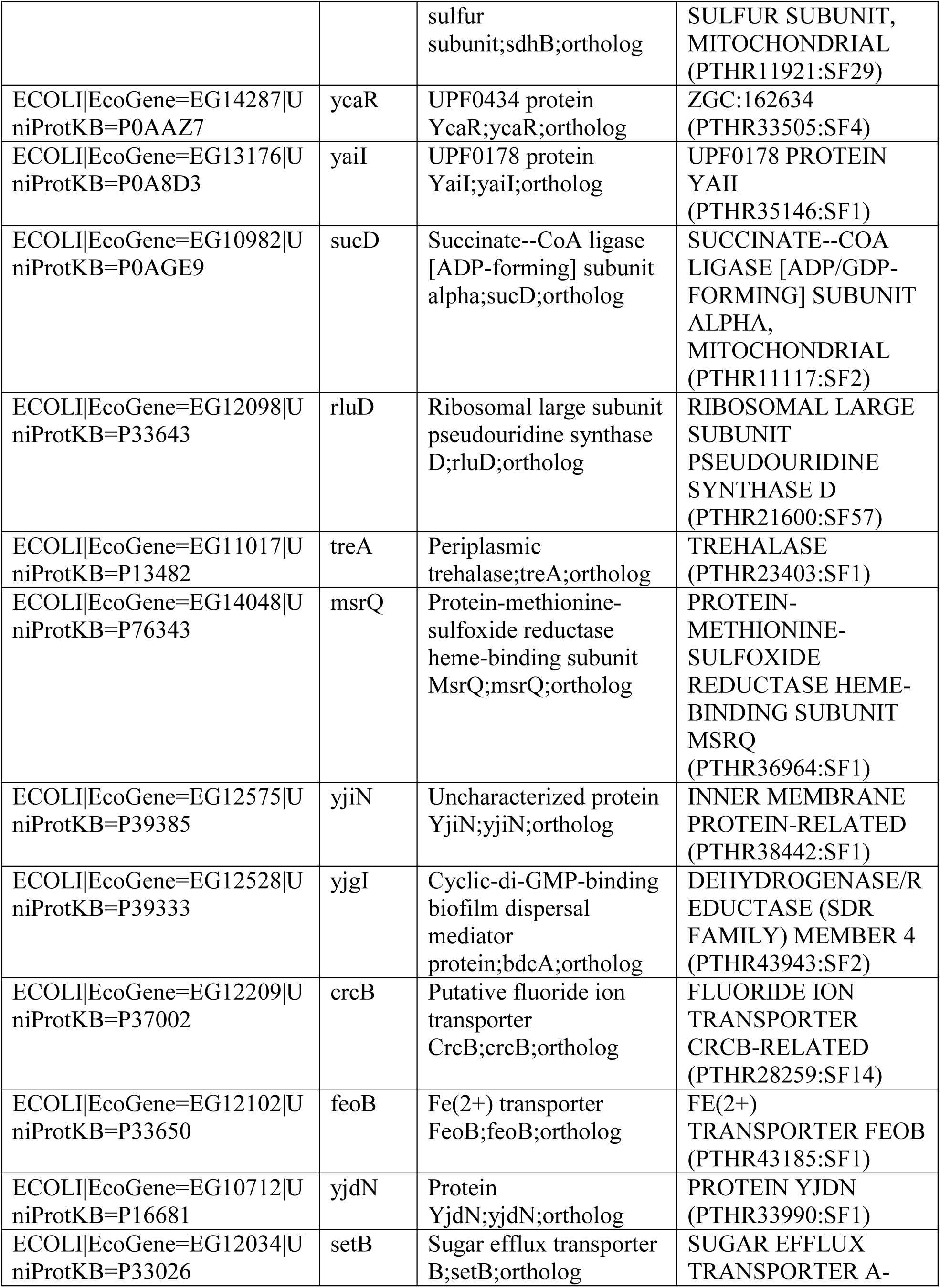
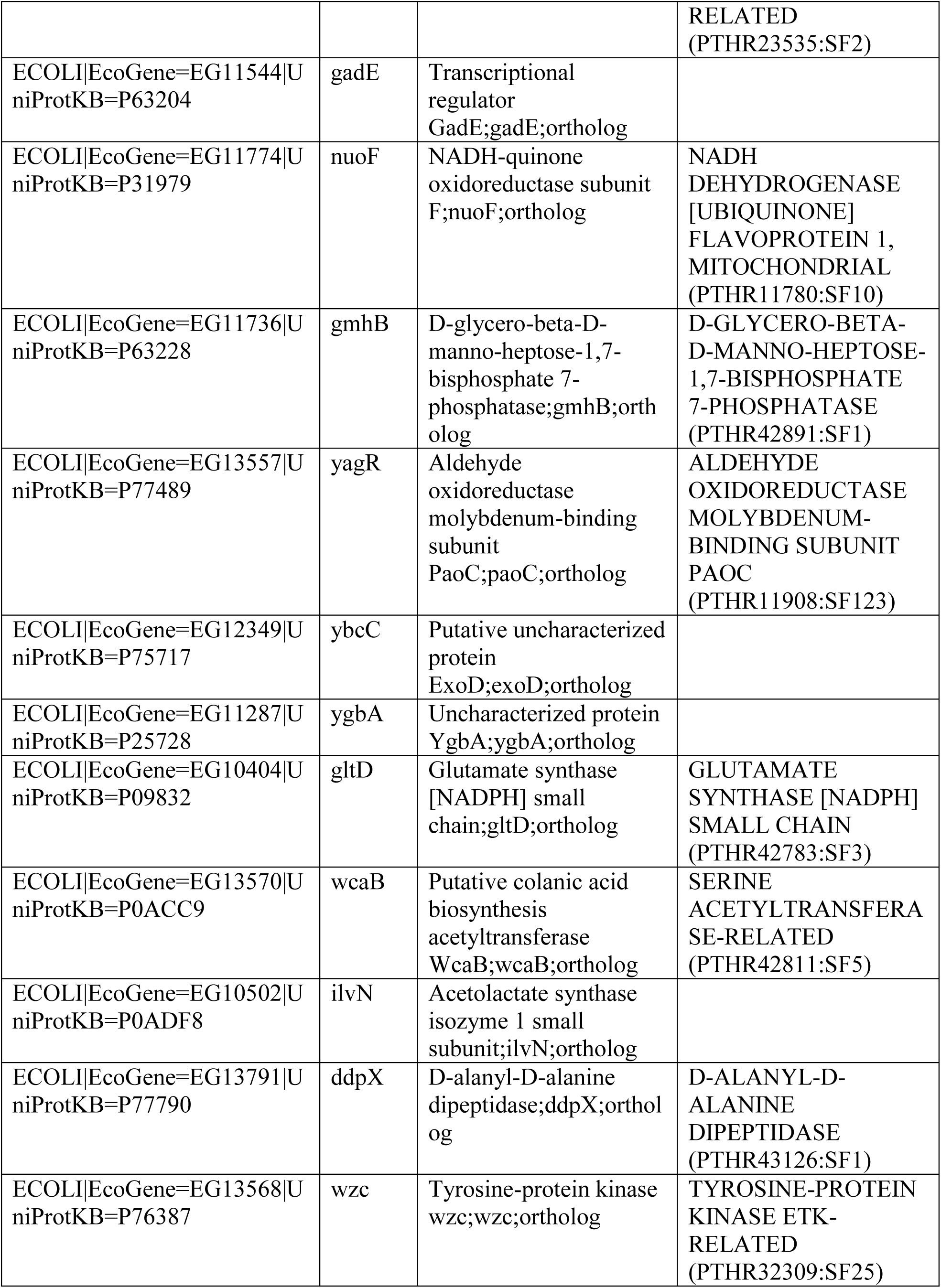
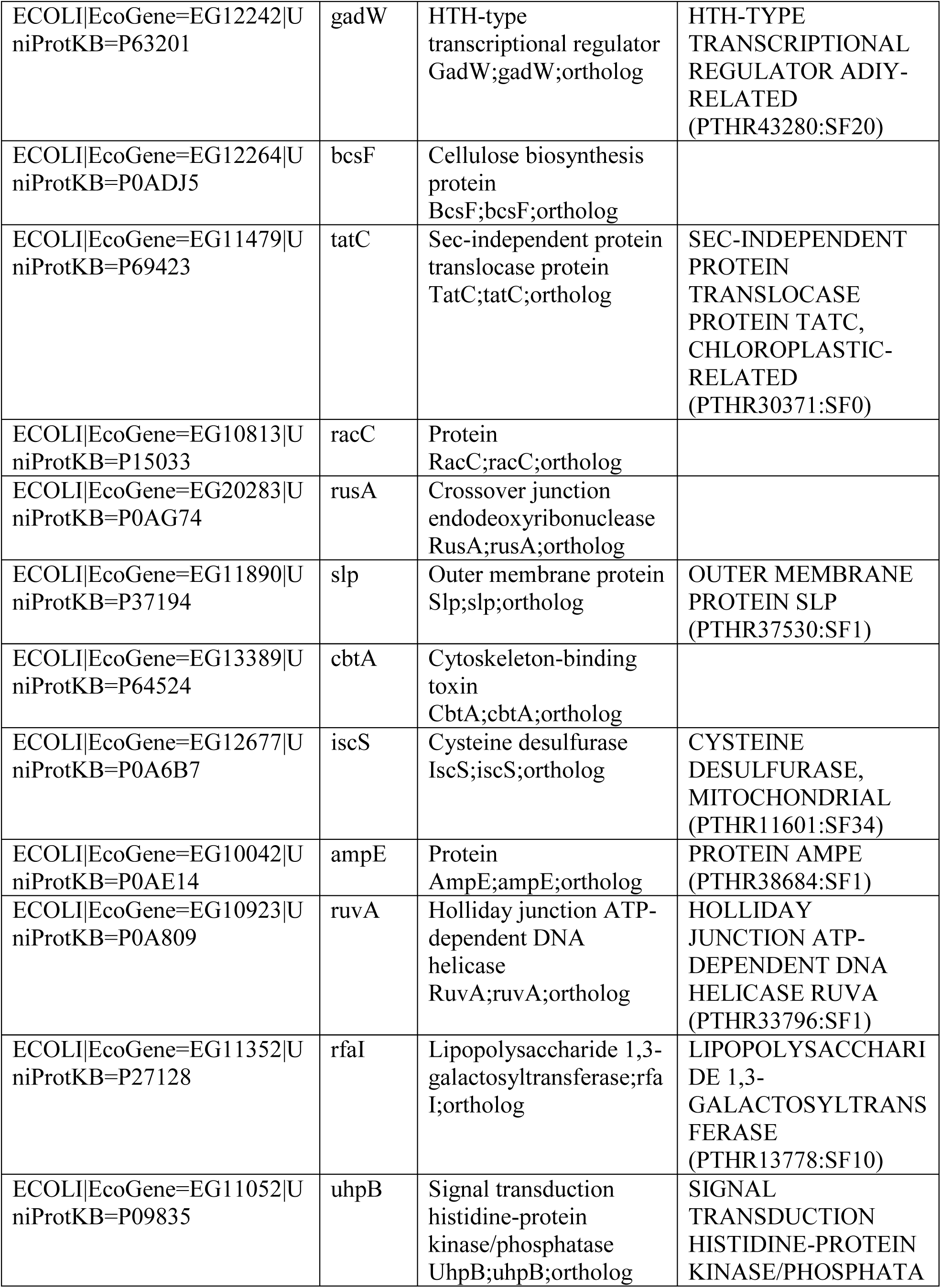
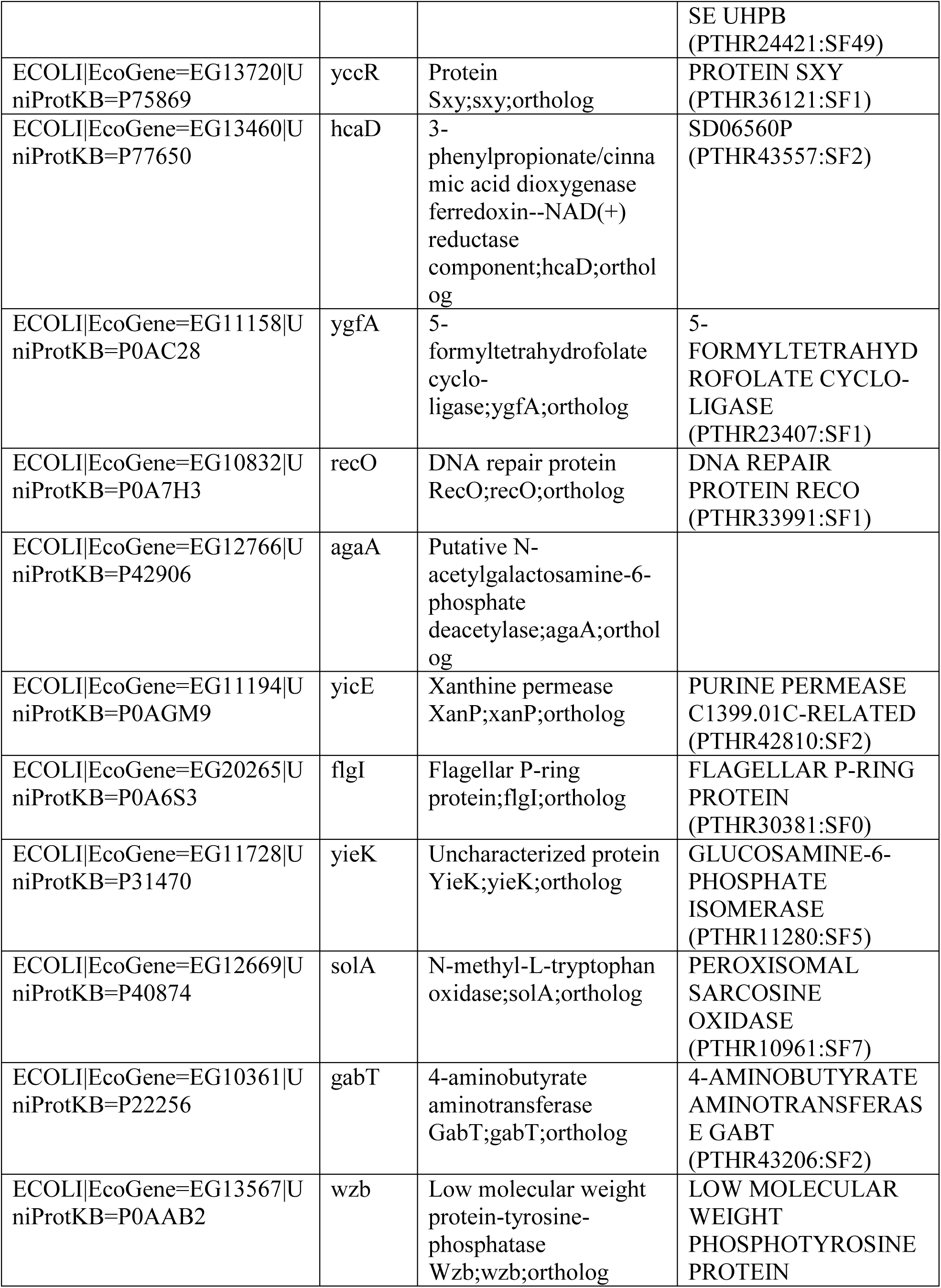
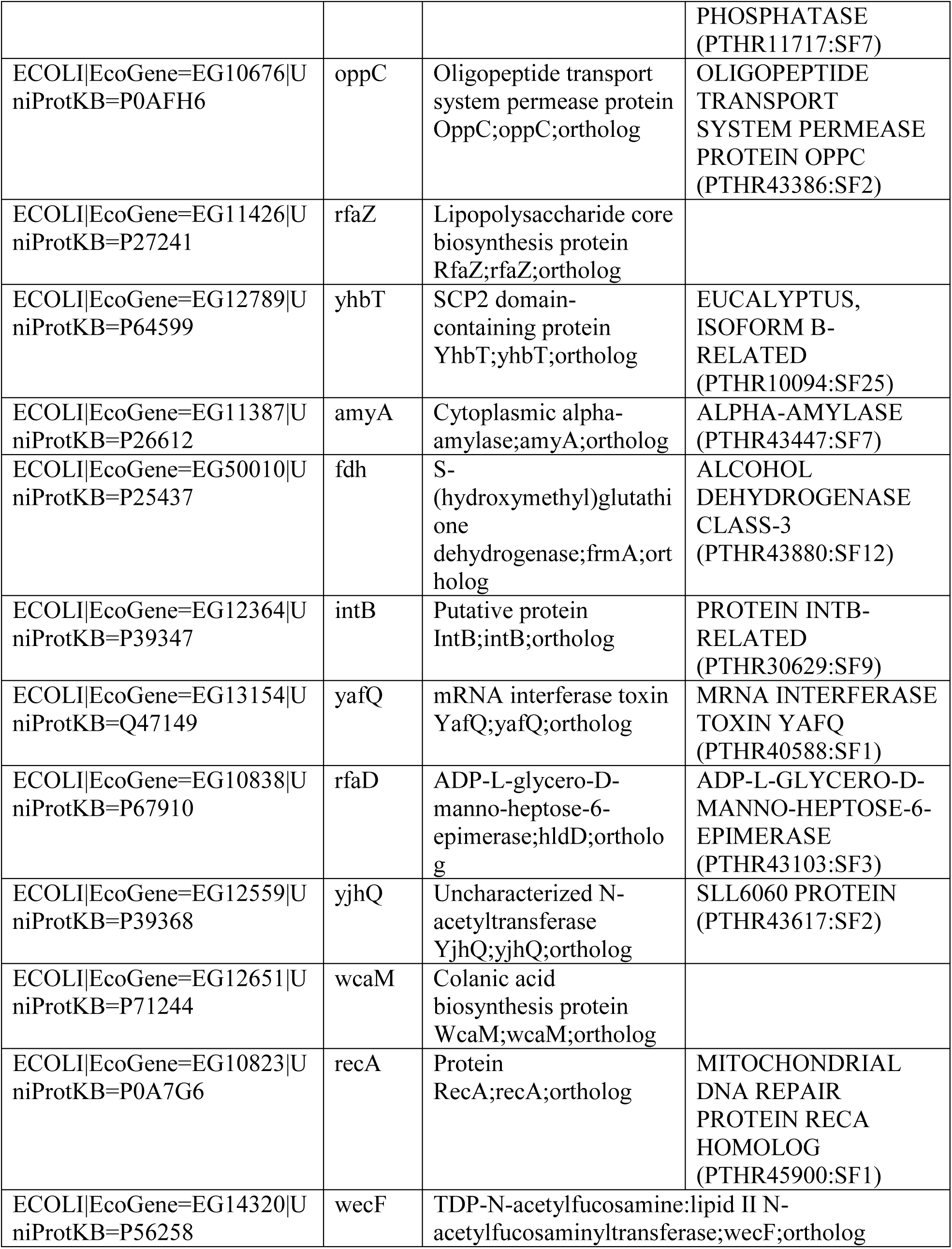
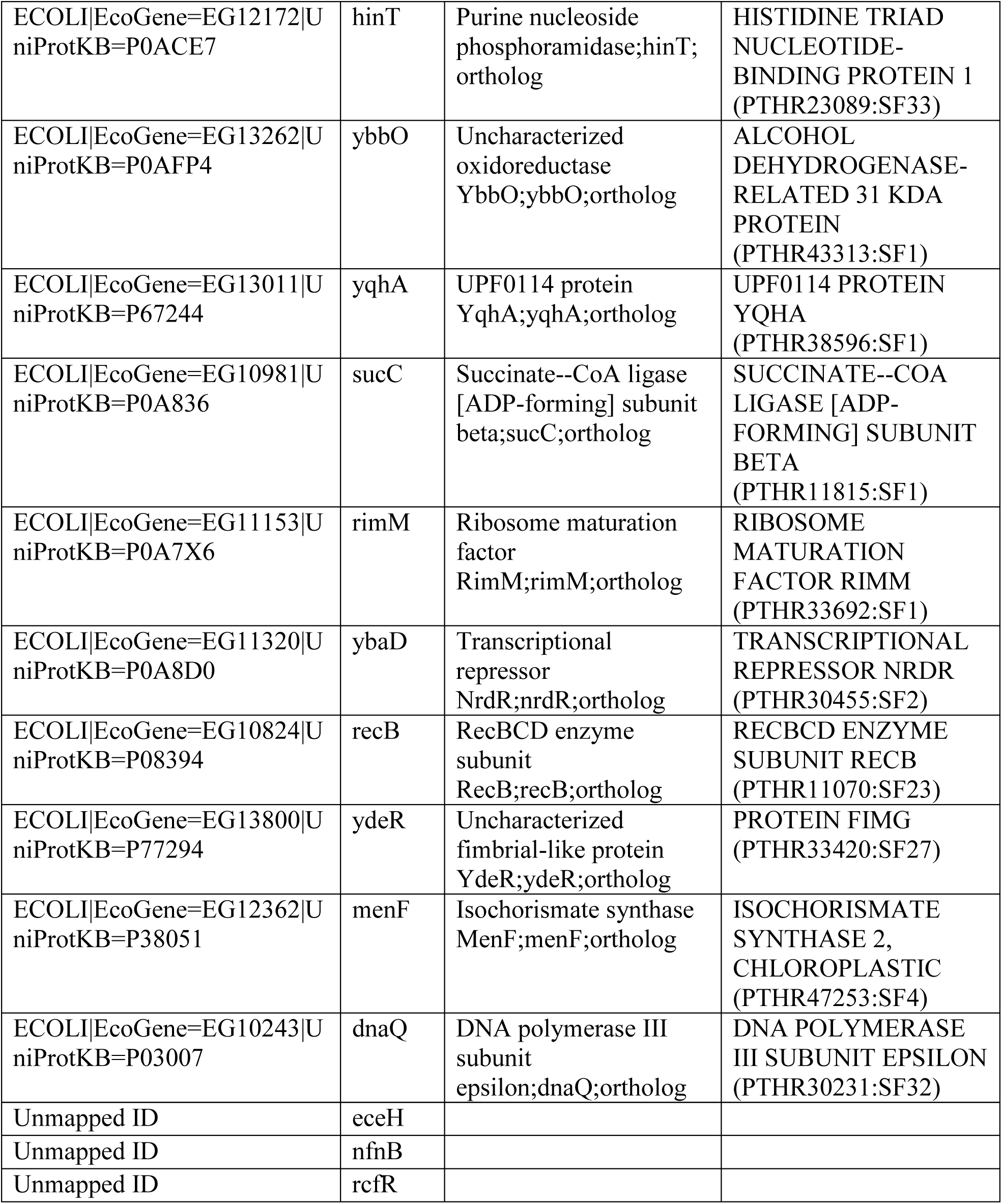
List of genes knockouts that reduced calcium AUC.

**Table S2:**
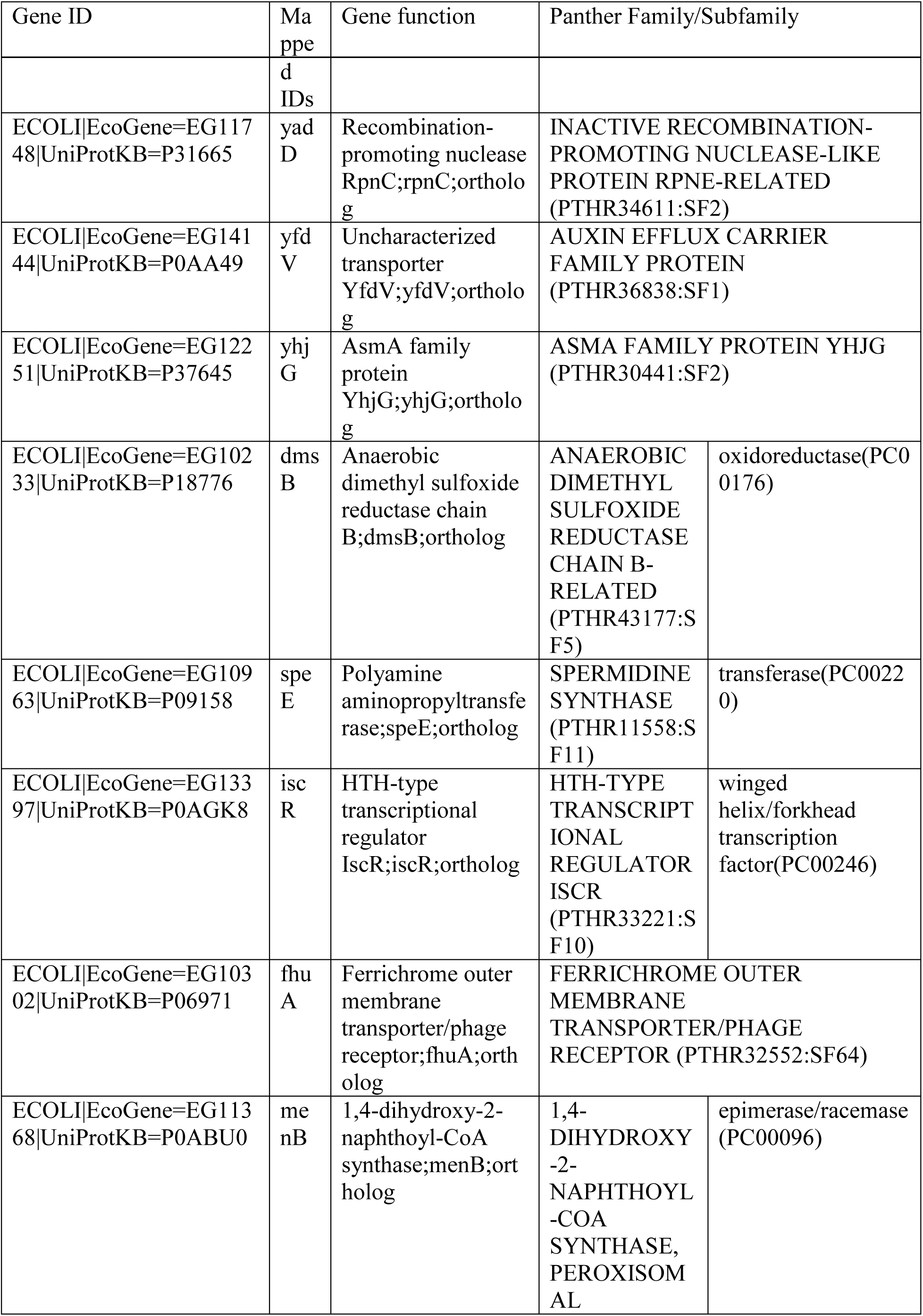
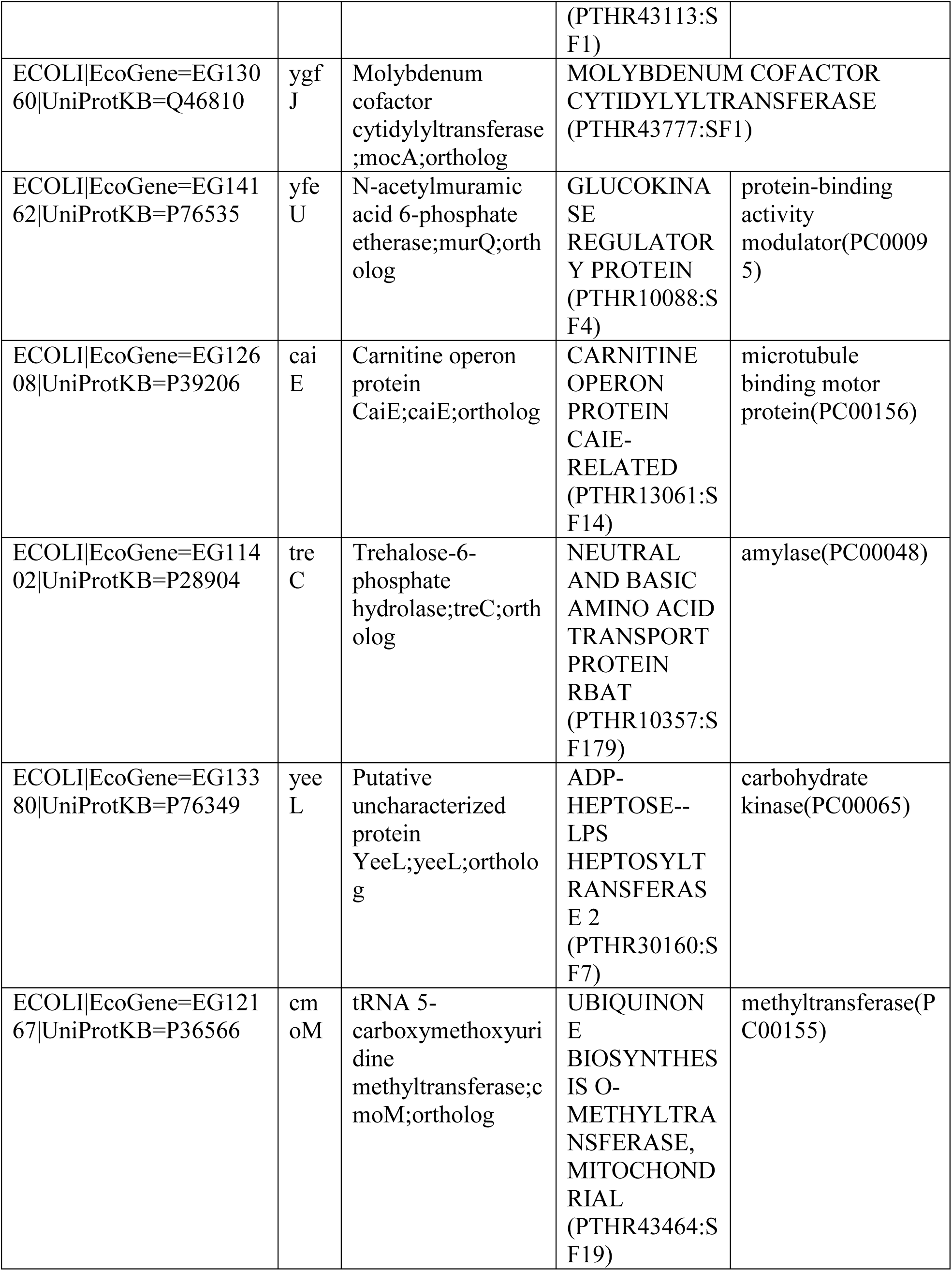
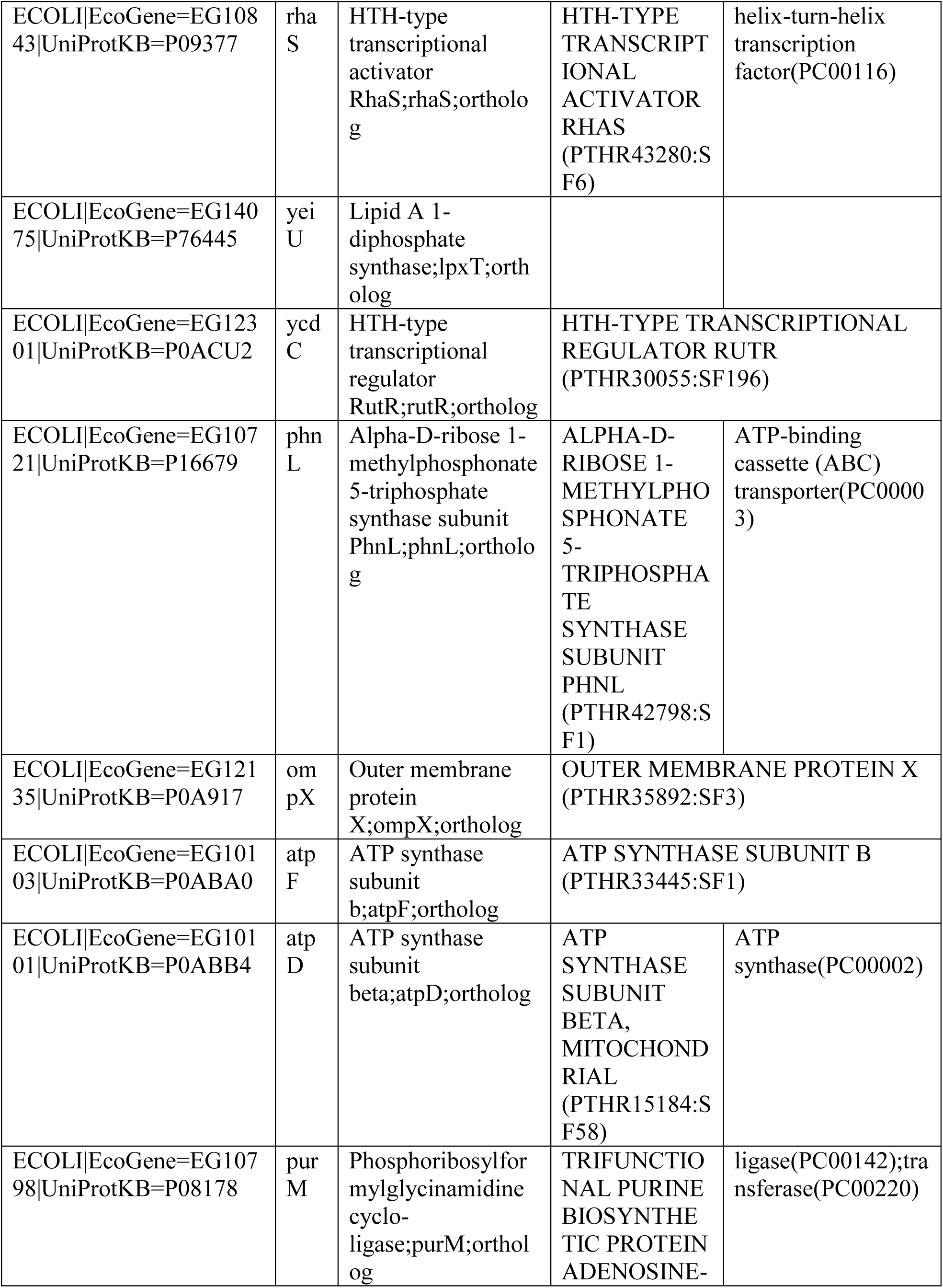
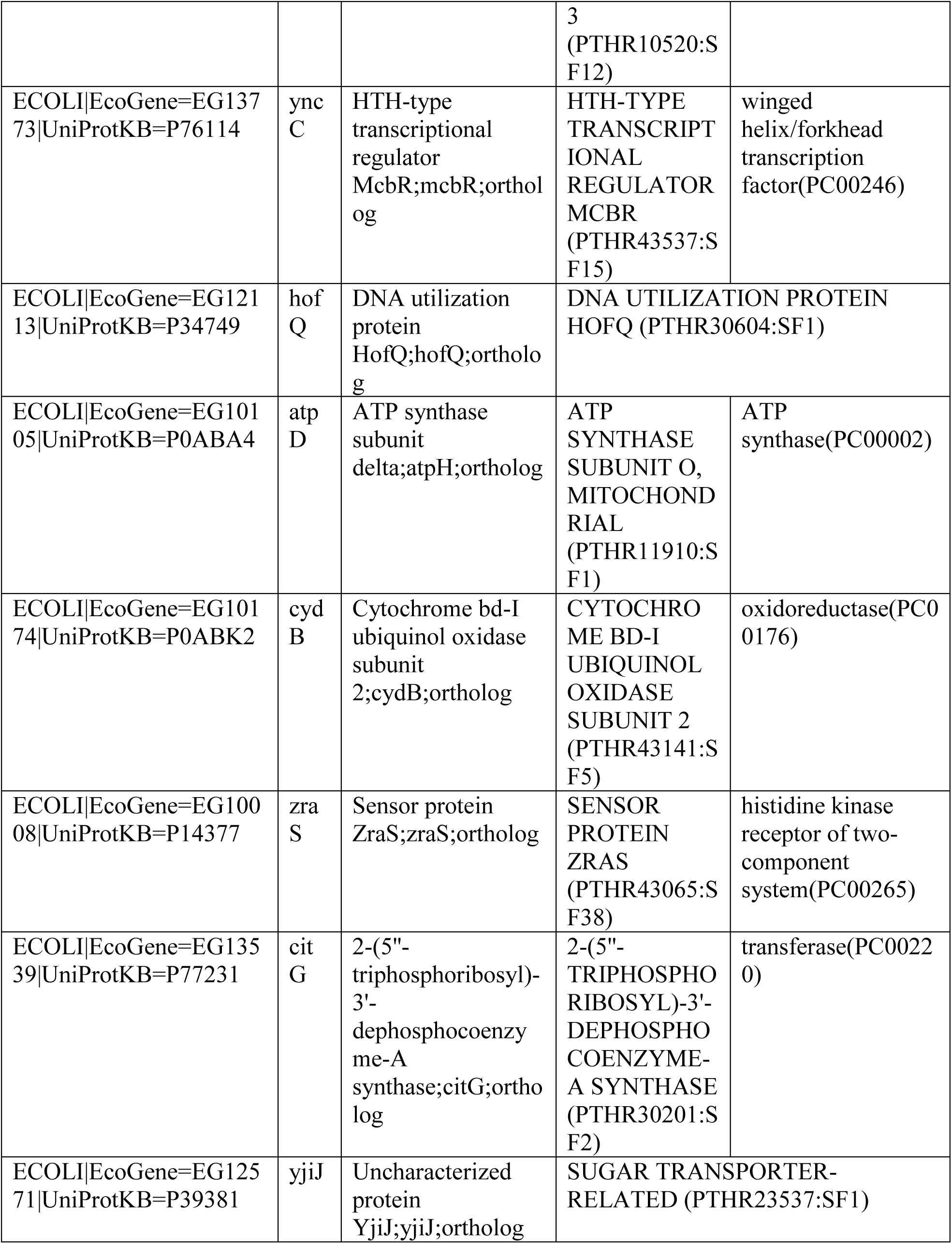
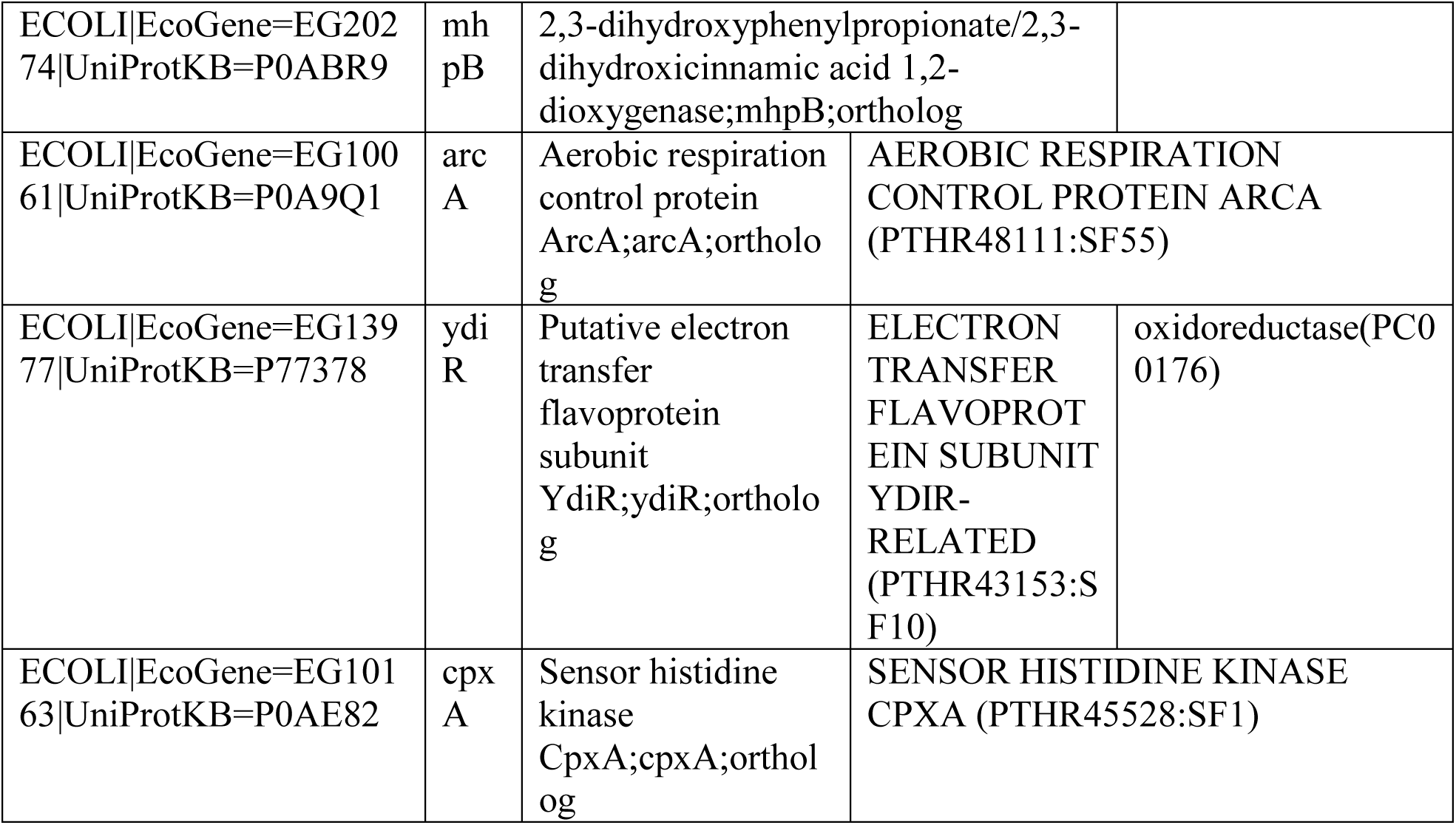
List of gene knockouts that increased calcium AUC.

